# Cell-to-cell diversification in ERBB-RAS-MAPK signal transduction that produces cell-type specific growth factor responses

**DOI:** 10.1101/704429

**Authors:** Hiraku Miyagi, Michio Hiroshima, Yasushi Sako

## Abstract

Growth factors regulate cell fates, including their proliferation, differentiation, survival, and death, according to the cell type. Even when the response to a specific growth factor is deterministic for collective cell behavior, significant levels of fluctuation are often observed between single cells. Statistical analyses of single-cell responses provide insights into the mechanism of cell fate decisions but very little is known about the distributions of the internal states of cells responding to growth factors. Using multi-color immunofluorescent staining, we have here detected the phosphorylation of seven elements in the early response of the ERBB–RAS–MAPK system to two growth factors. Among these seven elements, five were analyzed simultaneously in distinct combinations in the same single cells. Although principle component analysis suggested cell-type and input specific phosphorylation patterns, cell-to-cell fluctuation was large. Mutual information analysis suggested that cells use multitrack (bush-like) signal transduction pathways under conditions in which clear cell fate changes have been reported. The clustering of single-cell response patterns indicated that the fate change in a cell population correlates with the large entropy of the response, suggesting a bet-hedging strategy is used in decision making. A comparison of true and randomized datasets further indicated that this large variation is not produced by simple reaction noise, but is defined by the properties of the signal-processing network.

**Author Summary:** How extracellular signals, such as growth factors (GFs), induce fate changes in biological cells is still not fully understood. Some GFs induce cell proliferation and others induce differentiation by stimulating a common reaction network. Although the response to each GF is reproducible for a cell population, not all single cells respond similarly. The question that arises is whether a certain GF conducts all the responding cells in the same direction during a fate change, or if it initially stimulates a variety of behaviors among single cells, from which the cells that move in the appropriate direction are later selected. Our current statistical analysis of single-cell responses suggests that the latter process, which is called a bet-hedging mechanism is plausible. The complex pathways of signal transmission seem to be responsible for this bet-hedging.

## Introduction

Cellular responses to environmental changes occurring inside and outside cells are regulated by intracellular reaction networks for signal processing that contain various levels of fluctuation. The same reaction network is sometimes used to process multiple types of input signal, and the response to the same signal is dependent on the cell type and the cellular environment. Moreover, cellular information processing often includes large cell-to-cell variance [1–5] even within the same cell type under the same conditions. Signal processing networks usually regulate a variety of elemental cellular reactions, including morphological changes, migration, gene expression, and metabolism, and variations in the combination of these multiple reactions result in cell-to-cell fluctuations in fate changes. However, very little is still known about these single-cell fluctuations in cell signaling.

Conversely, the distribution of single-cell responses to the same input signal can be analyzed statistically to provide information about the mechanisms of intracellular signal processing. The correlations between the responses of multiple elements in single cells are especially useful for quantifying the information flows along networks. For this purpose, the simultaneous responses of multiple elements must be measured in single cells. Although there have been several theoretical studies of the correlation of multiple elements in the same networks [6–9], there have been few experimental studies in the field of signal transduction [4, 10, 11] when compared to genomics and proteomics [12, 13].

In our present study, we analyzed the ERBB–RAS–MAPK system responsible for cell fate decisions, including proliferation, differentiation, survival, and death, in various types of animal cells [14]. The dysregulation of this system results in genetic diseases and carcinogenesis. A family of receptor tyrosine kinases, designated ‘ERBB’, on the plasma membrane binds various types of growth factors (GFs) to activate this system [15]. ERBB consists of four members, ERBB1 (epidermal growth factor receptor) to ERBB4, which display specific ligand selectivity as homo- and heterodimers. The activation of the ERBBs upon ligand association stimulates multiple signaling pathways inside cells, including the calcium response pathway and the RAS and AKT activation pathways. Variations in the combinatorial activation of these pathways must be associated with the multimodal functions of the system, but the details remain largely unknown.

To date, more than 100 different elements of the ERBB–RAS–MAPK system have been identified [16]. We selected seven of these in our current study and detected their activation in single cells: four ERBBs, Ca^2+^/calmodulin-dependent protein kinase II (CaMKII), phosphatidylinositol 4,5-bisphosphate (PIP2), and extracellular signal-regulated kinase (ERK). Cells were stimulated with epidermal growth factor (EGF) or heregulin (HRG) to induce the activation of the ERBBs. EGF and HRG are the direct ligands for ERBB1 and ERBB3/B4, respectively [15]. The association of these GFs induces tyrosine phosphorylation in the cytoplasmic tails of the ERBBs. Not only the direct receptors, but other ERBBs are also phosphorylated through their heterodimerization with the direct receptors. CaMKII is autophosphorylated and activated when it associates with Ca^2**+**^**/**calmodulin [17]. ERBB phosphorylation activates the IP3 pathway through the recruitment of phospholipase Cγ (PLCγ) to the plasma membrane to increase the cytoplasmic Ca^2+^ concentration, which increases the Ca^2+^/calmodulin level and results in CaMKII phosphorylation. The density of PIP2 in the plasma membrane is also affected by ERBB signaling [14]. In addition to the recruitment of PLCγ to digest PIP2, generating IP3, the recruitment of phosphoinositide 3-kinase by the activated ERBBs and RAS converts PIP2 into phosphatidylinositol 3,4,5-trisphosphate (PIP3). PIP3 production is a signal to the AKT pathway. ERK is a kind of mitogen-activated protein kinase (MAPK) that acts downstream in the RAS pathway. The small G protein, RAS, is activated by a guanine nucleotide exchanger, son of sevenless (SOS). SOS forms a complex with GRB2 and is recruited to the plasma membrane through the association between phosphorylated ERBBs and GRB2, to activate RAS. Thus, CaMKII, PIP2, and ERK are elements operating downstream from the ERBBs in the ERBB–RAS–MAPK system. The pathways from the ERBBs to these three elements are distinct but interconnected. After phosphorylation, the ERBBs show different affinities for cytoplasmic proteins connecting to the distinct pathways [18]. However, the heterodimerization and transactivation of the ERBBs and the interconnections (cross-talk) between their downstream pathways make the situation complex.

The cell-type specific response of the ERBB–RAS–MAPK pathway is not yet fully understood in relation to GF specificity. For example, in the human-cancer-derived cell line MCF7, EGF and HRG stimulate cell proliferation and differentiation, respectively [19]. As mentioned above, EGF and HRG associate with different types of ERBBs. However, considering the heterodimerization among the ERBBs and the commonality in the downstream pathways for each ERBB, it is not easy to imagine the mechanism underlying this GF specificity. Based on the collective responses of cell populations, EGF and HRG are called proliferation and differentiation factors, respectively. However, in PC12 cells, in which EGF and nerve growth factor (NGF) act as proliferation and differentiation factors, respectively, we previously found that both EGF and NGF increase the rates of proliferation and differentiation simultaneously [5]. According to our prior analysis, the differences in the functions of EGF and NGF are caused by the irreversibility of the differentiation that occurs in the presence of NGF. In this case, the specificity of the cell fate change is a population behavior, and not deterministic in each single cell.

In our present study, we compared three human epithelial carcinoma cell lines, HeLa, A431, and MCF7, during stimulation with EGF or HRG. The cellular response to these GFs at the population level is specific to each cell type: HeLa and MCF7 cells are stimulated to proliferate in the presence of EGF, but A431 shows growth arrest, at least at nanomolar concentrations of EGF [20, 21]. MCF7 cells differentiate to acquire mammalian gland cell-like properties in the presence of HRG [19], whereas HeLa and A431 cells show controversial or obscure responses to HRG. We measured the responses of our seven selected elements in the ERBB–RAS–MAPK system in single cells stimulated with EGF or HRG, and analyzed them statistically. A correlation between the property of information transmission and the variety of response was detected, suggesting a cellular population strategy that induces fate change.

## Results

### Growth factor stimulation of the ERBB–RAS–MAPK system

The responses of the ERBB–RAS–MAPK system in three types of cultured cells derived from human epithelial carcinomas (HeLa, A431, and MCF7) to two types of GFs (EGF and HRG) were examined. EGF is a ligand for ERBB1 and HRG is a ligand for ERBB3 and ERBB4. Both GFs activated the ERBB–RAS–MAPK system in all three types of cells, which was detected as increases in the phosphorylation levels of the ERBBs and ERK (Fig. S1). The concentrations of 20 nM EGF and 30 nM HRG, which caused almost saturated responses in all three cell types after stimulation for 5 min, were used in the subsequent experiments. In our current analyses, we focused on the early stages (5 min) of signal transduction to examine the direct effects of ERBB activation on the downstream molecules, minimizing the effects of the signal termination of the ERBBs and the long negative feedback from ERK.

### Multicolor immunofluorescence detection of single-cell responses

The phosphorylation levels of two types of ERBBs (ERBB1 and one of ERBB2–B4), CaMKII, and ERK were detected simultaneously with PIP2 in single cells using multicolor immunofluorescence staining (Fig. S2 and Table S1). In addition to the images of the signaling elements acquired in five channels, an additional image of the autofluorescence (AF) channel was acquired under deep blue excitation for later spectral unmixing. Based on the emission spectra of the fluorescent dyes and AF, the signals in individual channels were successfully unmixed (Fig. S3), after which the fluorescence signals were averaged over the entire area of a single cell so that no subcellular spatial information was considered. We analyzed isolated cells to minimize the effects of cell-to-cell adhesion.

The response patterns of the signaling molecules were characteristic of the cell type and the GF (Figs 1 and S4), as was clearly observed in the phosphorylation of the ERBBs and ERK. In some cases, especially for PIP2, negative responses (reduction in PIP2 density) were observed, as expected for the reaction network of the ERBB–RAS– MAPK system. On the whole, these results were reproducible in repeated independent experiments. However, the responses of some molecules, including ERBB2, CaMKII, and PIP2, fluctuated from time to time, suggesting large cell-to-cell variations (Fig. S4).

**Figure 1.**
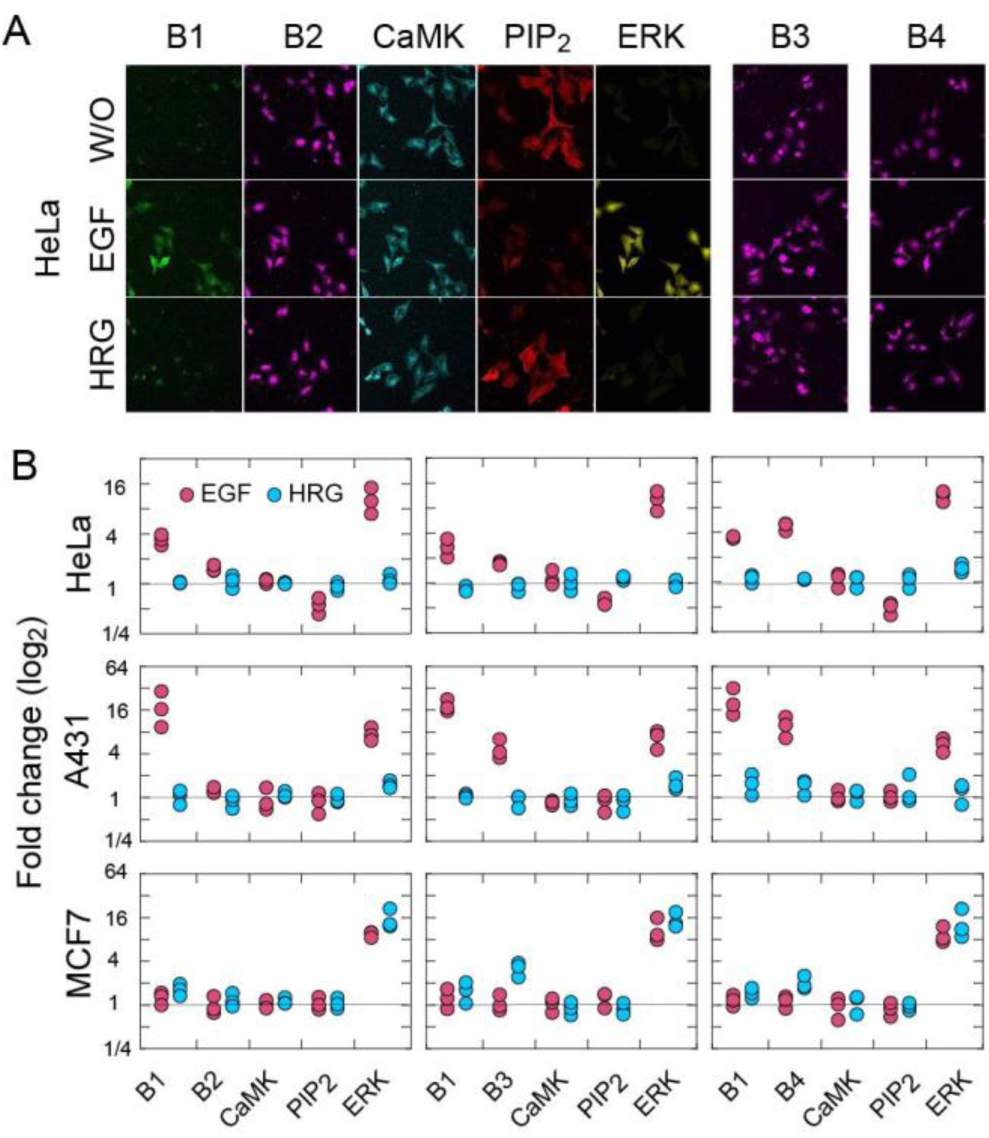
Immunofluorescence staining of cell signaling molecules. (A) Typical results for the multicolor staining of HeLa cells without (W/O) or with GF (EGF or HRG) stimulation. Fluorescence micrographs after spectral unmixing are shown. Images of pERBB3 (B3) and pERBB4 (B4) are the results of independent experiments from others. Field size, 325 × 325 μm^2^. (B) Fold changes in the fluorescence signals after GF stimulation plotted as averages of single cells. The number of cells measured was 115– 135 under each condition. The results of three independent experiments are shown. See Figure S4 for further details.

### Principle component analysis

The simultaneous detection of multiple molecular responses in single cells enables statistical correlation analysis to be performed. Figures 2 and S5 indicate the pairwise relationships between every combination of molecular responses in the same single cells. The noise from the immunostaining and detection experiments was small (9%–29% of the standard deviation [SD] of the signal distribution; Fig S6), so the distributions in Figures 2 and S5 predominantly reflect the distributions of the single-cell responses. The distributions were specific to the cell type and GF.

**Figure 2.**
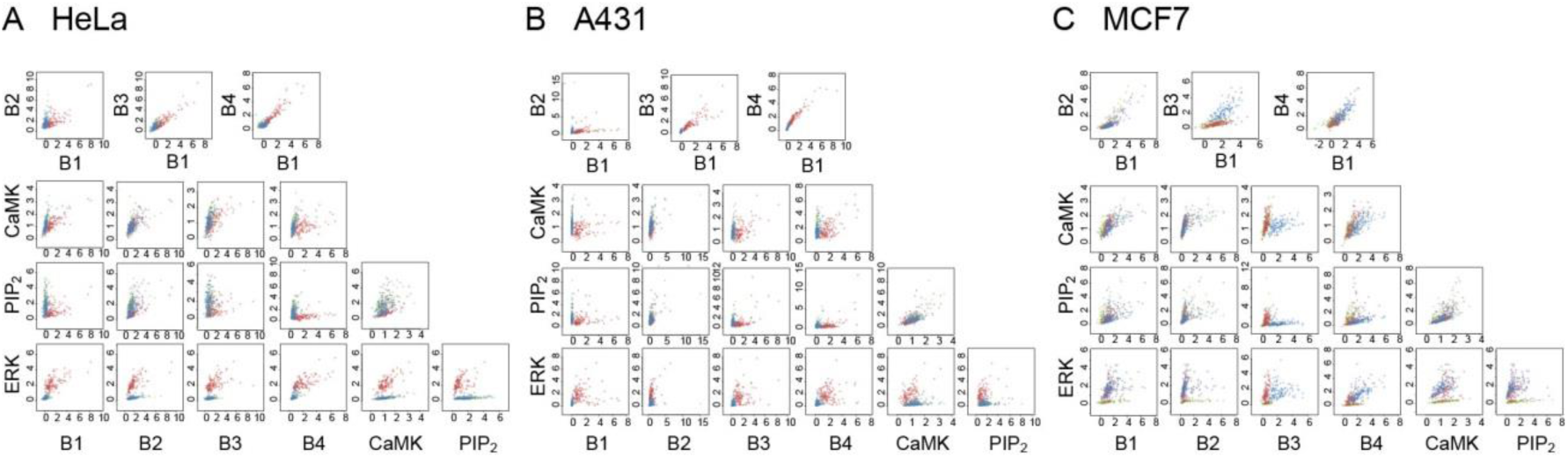
Two-dimensional distributions of single-cell staining intensities for HeLa (A), A431 (B), and MCF7 cells (C) without stimulation (green) and after stimulation with EGF (red) or HRG (blue). Intensities were normalized to the average for all the cells with and without stimulation. Typical results of three independent experiments are shown (see Figure S5 for the results of all experiments).

Principal component analysis (PCA) of the five channel molecular responses was carried out to detect cell-type and GF specificity. Under every condition, more than 73% of the single-cell distribution could be explained by the PC1 and PC2 scores (Fig. 3A). These scores indicate the deviations of a single-cell response along the orthogonal basis with the largest and the second largest variances in the five-dimensional single-cell distribution. While the PCA was independently performed for each cell type and ERBB combination, the profiles of the PC1 vector (rotation) under every condition indicated the positively correlated behaviors of all the elements (Fig. 3B). Under conditions in which a clear cell response to the GFs has been reported (HeLa and A431 with EGF, and MCF7 with EGF and HRG), the PC1 scores were relatively higher than those of PC2. On the other hand, under conditions showing an obscure cell response (HeLa and A431 with HRG), the PC2 scores were relatively higher. In the PC2 rotations of these cases, the ERBBs and ERK showed a negative correlation with CaMKII and PIP2. The proliferation (EGF) and differentiation (HRG) response patterns in MCF7 cells were distinct in terms of the PC2 scores. In the PC2 rotations of MCF7, the ERBB responses were negatively correlated with those of CaMKII and PIP2, as shown in other cells. However, the negative correlation between ERBB1/B2 and ERK was specifically observed in MCF7 cells.

**Figure 3.**
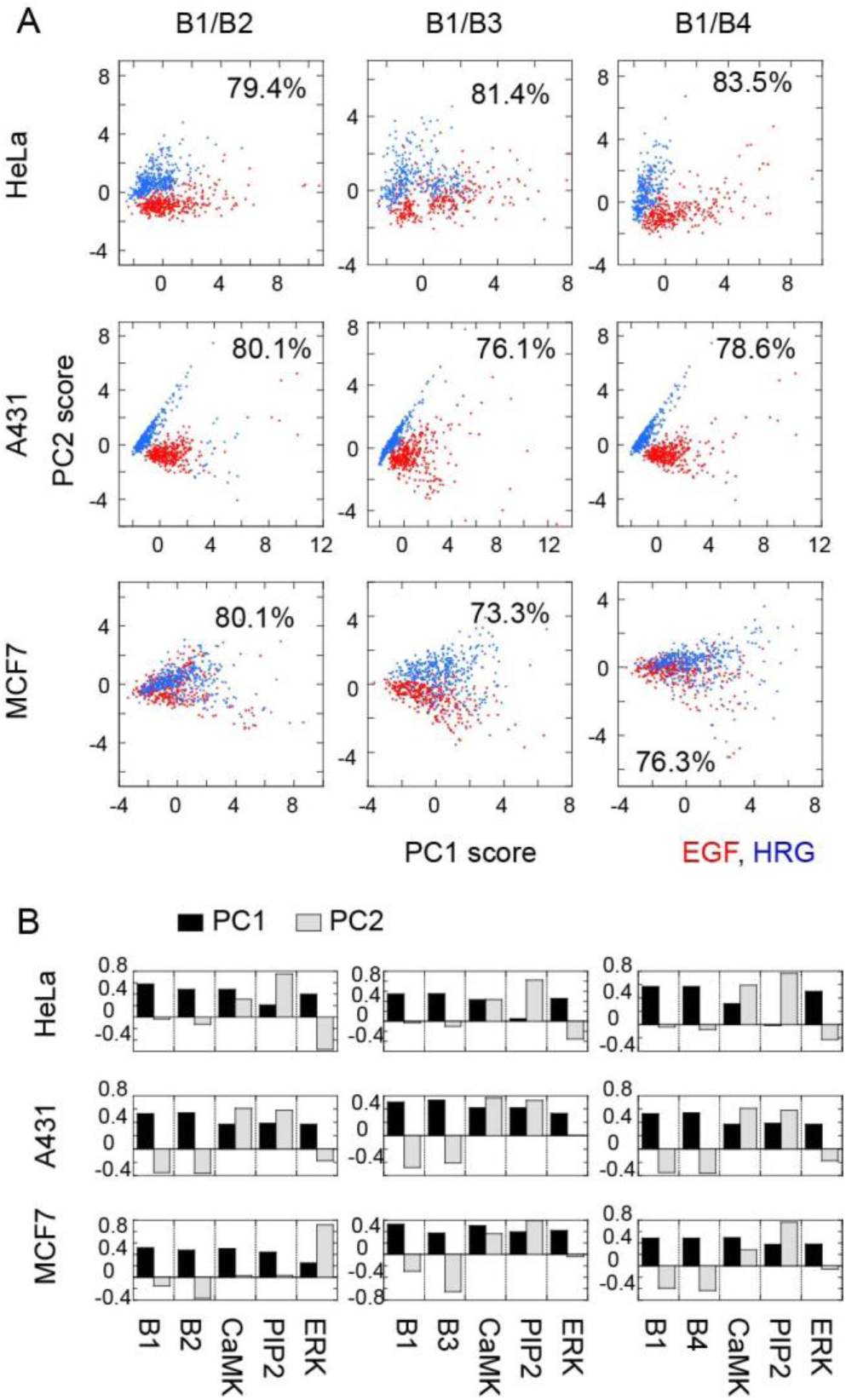
PCA of the single-cell response patterns. (A) Single-cell distributions in the PC1/PC2 space after EGF (red) and HRG (blue) stimulations. Percentage indicates the summation of the contributions of PC1 and PC2 to the whole distribution. (B) Rotations of PC1 (black) and PC2 (gray). PCA was performed independently for each cell type and combination of ERBBs, but conjointly between the two GFs.

### Mutual information analysis

In many instances, the distribution of single cell responses was found to be widely spread with low linearity, although almost all one-dimensional distributions showed a simple single peak (Fig. S5). Correlation coefficient analysis is inadequate for characterizing such distributions so mutual information (MI) was instead obtained as a measure of the pathway strength between two molecules (Figs S7 and 4A, Table S2). The MI is the amount of information about the response of one of the two molecules that can be obtained from the response of the other molecule, which is an extension of the correlation coefficient to non-linear relationships. The standard errors in the MI values were usually about 10% or less of the average values over the independent experiments, showing good reproducibility (Table S2).

In general, the MI between the ERBBs was larger than between the ERBBs and their downstream molecules (CaMKII, PIP2, ERK), as expected from the direct heterodimerization that occurs between the ERBBs. Of the pathways from the ERBBs to the three downstream molecules examined, those to CaMKII and PIP2 carried the most and least information, respectively, in most cases. CaMKII, to which the MI from the ERBBs was generally large, showed small fold changes in phosphorylation. It must be noted that the response intensity of the molecules, for example shown as the fold change in their average phosphorylation levels, do not necessarily correlate with the amount of MI (Fig. S8). Even when the level of phosphorylation is low, the amount of MI would be large if the downstream molecule read the upstream activation precisely. In A431 cells, however, both the phosphorylation levels of the downstream molecules and MI were generally small, as expected intuitively. Unexpectedly, the direct receptors of each GF (ERBB1 for EGF and ERBB3/B4 for HRG) did not always transmit a larger MI amount than the indirect ERBB families between the downstream molecules. This phenomenon suggests complex signal processing and/or pathway-specific noises in the network between the ERBBs and their downstream molecules.

The MI diagrams shown in Figure 4A indicate the thicknesses of the information transmission pathways under each condition. To characterize the specificity of the signal transduction network, the major pathways of information transmission were extracted from each diagram (Fig. 4B). As expected from the dense dimerization network among the ERBBs, we observed multiple pathways to their downstream molecules except in the A431 cells stimulated with HRG, in which ERBB2 was the privileged receptor molecule in transmitting information to the downstream molecules. Even under stimulation with EGF, the A431 cells used a smaller number of pathways for information transmission than the other cell types. In HeLa and A431 cells, the numbers of major pathways were larger during EGF signaling than during HRG signaling. Although the threshold level of MI (0.6 bit) used in Figure 4B was determined arbitrarily to capture the characteristic differences between the experimental conditions, independent of the threshold level, in the HeLa and A431 cells, EGF tended to stimulate larger numbers of information pathways than HRG (Fig. 4C). In contrast, EGF and HRG stimulated similar numbers of pathways in MCF7 cells. Thus, interestingly, larger numbers of pathways were used under conditions known to cause clear cell fate changes, such as proliferation, growth arrest, and differentiation, in response to GFs [19–21].

**Figure 4.**
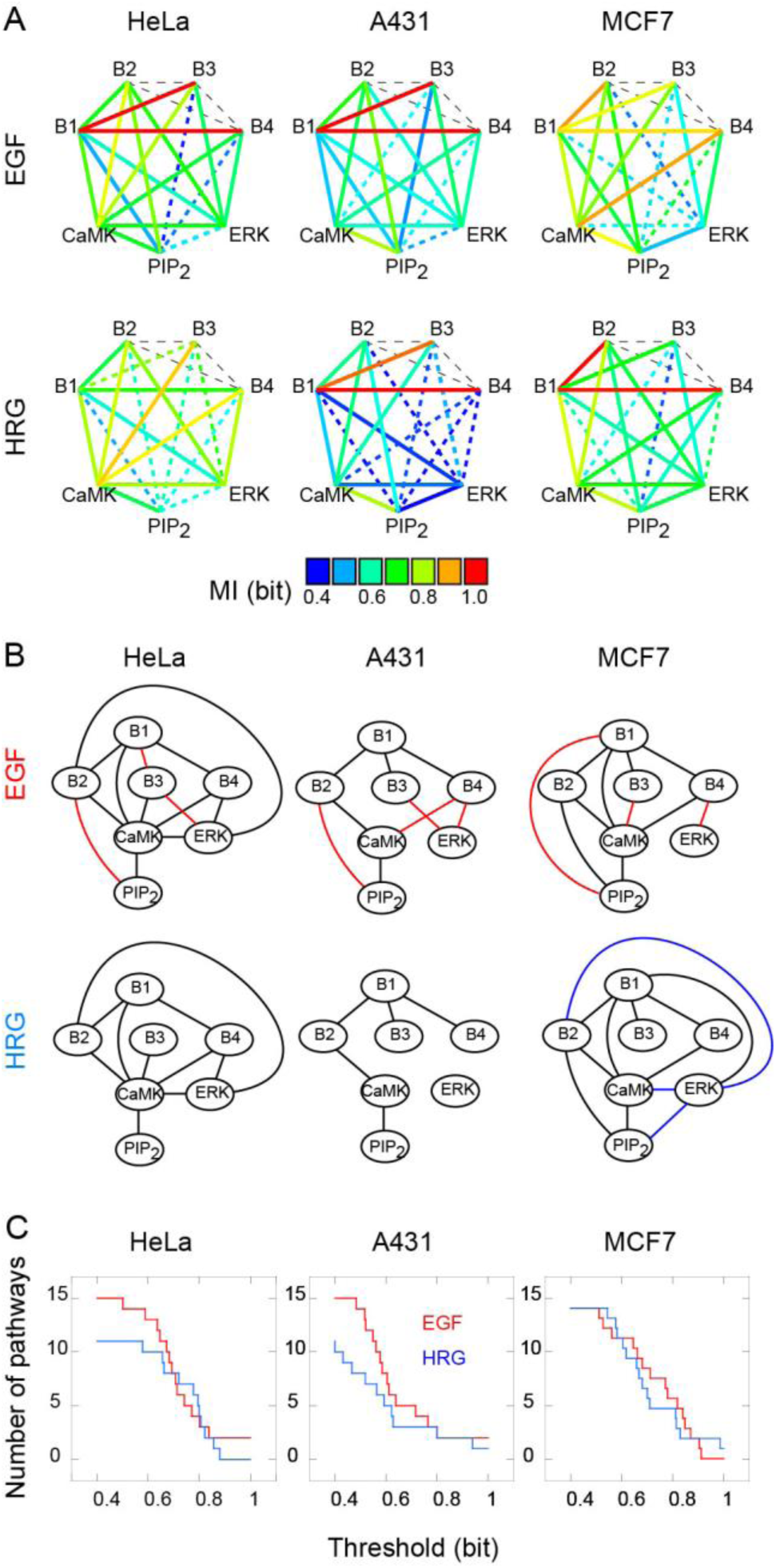
Mutual information (MI) analysis. (A) MI diagrams (see Table S2 for the average values and SE). Statistically insignificant pathways (P > 0.05 on *t* test) are indicated by the dotted lines. MI was not measured in the black dashed lines. (B) Networks of major information transmission pathways (MI > 0.6 bit and P < 0.05). Note that some other pathways carry smaller but significant amounts of information. Pathways specific for EGF and HRG in each cell type are colored red and blue, respectively. (C) Numbers of statistically significant pathways plotted as a function of the threshold MI values.

### Clustering of single-cell response patterns

The variations in single-cell responses were evaluated as the numbers of clusters in the five-dimensional molecular phosphorylation patterns, applying a hierarchical clustering method. In this method, the number of clusters depends on the threshold distance to distinguish individual clusters, and the decay curves in the cluster number as the increase of the threshold distance reflects the properties of the single-cell response distribution (Fig. 5A). In HeLa and A431 cells, the number of clusters was larger under EGF stimulation than under HRG stimulation, independent of the threshold value. Thus, the response variation was larger under conditions in which GF-induced cell fate changes have been reported. This clustering property remained after the dataset was randomized by permutation (Fig. 5B). One clear difference between the true and randomized datasets was that the decay in the randomized data displayed sharp cut-offs in the large threshold tails, indicating that the dynamic ranges of the true cell responses in multidimensional space were wider than expected from the random responses. Comparison of the variance in the single-cell responses between the true and randomized data (Fig. 5C) also indicated larger variations in the true cell responses under conditions that induce clear cell fate changes.

**Figure 5.**
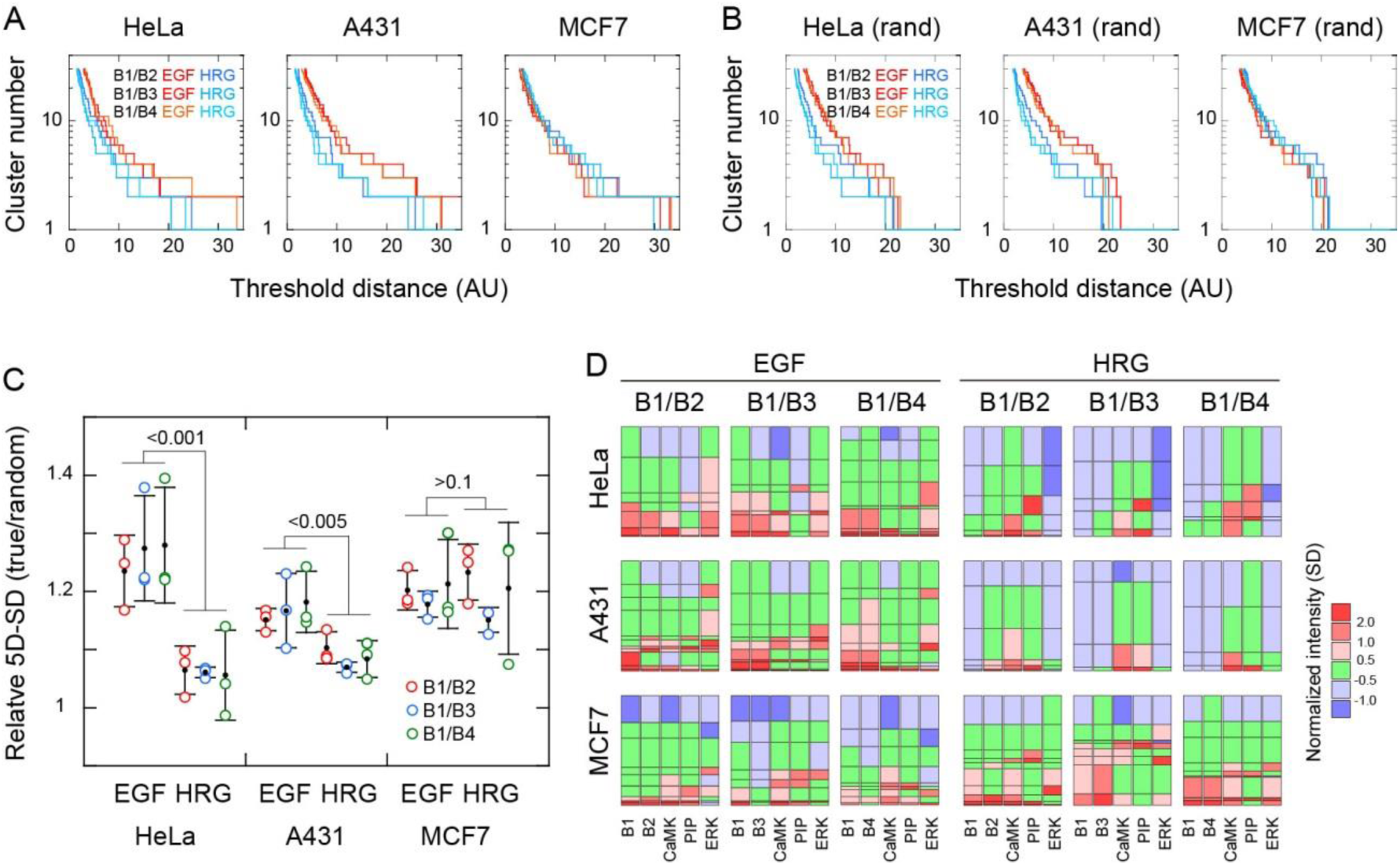
Variations in single cell responses. (A, B) Single-cell response patterns were classified using Ward’s method. Numbers of clusters are shown as a function of the threshold distance to define the clusters in the true (A) and randomized datasets (B). AU, arbitrary unit. (C) Ratios of five-dimensional SD (5D-SD, defined as (∑_*i*_ SD_*i*_^2^)^1/2^) for the true and randomized data were plotted. SDi is the SD of the normalized single-cell responses of each molecule. Significance levels on one-way ANOVA are shown. (D) Results of clustering with a threshold distance of 8. Average responses of each cluster are shown in six-step color cords. The height of each rectangle indicates the fraction of cells belonging to each cluster. The average and SD of each cluster is listed in Table S3.

To see the specific response patterns, Figure 5D was generated at a threshold distance that was determined arbitrarily but based on the distance/cluster number relationships shown in Figure 5A to capture the characteristics of each condition. In the results, the clusters aligned as the intensities of the ERBB responses increased (see Table S3 for the intensity values in each cluster). Unexpectedly, the intensities of the downstream responses did not correlate monotonically with those of the ERBBs in many cases. Instead, in each cell type, similar ERBB responses induced multiple distinct patterns of downstream responses, and strong downstream responses were observed under the medium intensities of the ERBB responses.

Shannon’s entropy in the cluster distribution of the cell responses is another indication of the variation in the cell-to-cell responses. We found that these entropy values also correlated with the collective cellular responses to GFs, insofar as large entropy values were observed under conditions that induced cell fate changes in the population (Fig. S9A). The small entropy values in HeLa and A431 cells stimulated with HRG might be attributable to the weak activation of some elements, which cannot be easily divided into numbers of clusters (Figs 1 and 2). However, strong activation in the averaged cells does not necessarily result in large entropy. Notably in our current analyses, the experimentally observed cell-to-cell variations were not simply produced by random noise, in which case, entropy size would increase with the average intensity. As expected from the significant amounts of MI (Fig. 4) in the regions of small and medium threshold distances, the entropy values for the true cell datasets were smaller than those after randomization, but larger in the regions of long threshold distances (Fig. S9B). This result indicates that the structured response distribution in true cells is a determining factor for the cell-to-cell variance.

## Discussion

Intracellular signal transduction networks consist of complicatedly connected reaction pathways involving a huge number of reaction elements. The ERBB–RAS–MAPK pathway is a typical example of such a cell signaling network. We here studied the origin and properties of the cell-to-cell deviations in the responses of this network to GF stimulation by observing the simultaneous activation of multiple elements in single cells. Although each element exhibited varied activation strength specifically depending on the GFs and cell type, it is not easy to quantitatively compare the importance of each elementary pathway or reaction in a diversified network. Because the function and activity of each are unique, a comparison based on the number or concentration of activated molecules, as detected with conventional biochemistry, is inadequate. One possible method of comparison is to measure the amount of information transmitted between molecular species, which allows quantification in a common unit i.e. the bit. Information analysis is a generalization of the linear correlation analysis for non-linear systems. To calculate the amount of information transmitted, the joint probability distribution of the responses is required. This means that the information analysis in a population of cells is essentially an analysis of the cell-to-cell deviations.

We measured multiple-element responses simultaneously in single cells. Seven elements were selected from the ERBB–RAS–MAPK system, including four receptors (ERBB1–B4) for GFs and three downstream molecules (CaMKII, PIP2, and ERK) in diverse but overlapping signaling pathways from these receptors. A theoretical analysis of the ERBB–RAS–MAPK system has suggested that these seven molecules are key factors determining the activation state of this network [22]. One remarkable property of the ERBB–RAS–MAPK system is that it regulates diverse cellular responses, including cellular proliferation, differentiation, survival, and movement, according to the cell type and input signal [15, 16]. From the statistics of the cellular responses, we gained insights into the mechanisms of these cell-type- and GF-specific responses and the cell-to-cell variations in them.

Three human epithelial-cancer-derived cell lines were used in this study. Each of these cell types shows distinct responses to specific GFs (EGF and HRG) via ERBBs. Upon stimulation with EGF, HeLa and MCF7 cells are induced to proliferate, whereas growth arrest is induced in A431 cells. Upon stimulation with HRG, MCF7 cells are induced to differentiate, whereas no clear effect has been observed previously in HeLa or A431 cells [19–21]. Interestingly, the connections between the major signal transduction pathways that involve the seven elements we measured herein were found to be more complex, containing multiple edges and loops (bush model; [23]) under conditions in which clear cell fate changes are induced by GFs than under other conditions (Fig. 4). The bush-like networks we observed indicate that various signal transmission pathways, or the activities of various elements, are used almost equally in population-level signal processing. Our PCA results also suggested a requirement for global activation of the network elements to induce cellular responses to GFs (Fig. 3). In contrast, under conditions in which the cellular responses to GFs were obscure, the major MI networks were simpler, with smaller numbers of significant edges (tree model; [23]). Information transmission in tree-like networks relies on monotonic signal transduction pathways in every cell. We found no clear relationship between the intensity of molecular activation and the amount of MI. Thus, MI analysis to detect correlations between molecular responses provides unique information about signal processing.

We found that the variation in single-cell responses, measured as the number of clusters (Fig. 5) and the entropy in the cluster distributions (Fig. S9) of single-cell responses, was large under conditions in which the collective cell fate changes have been reported previously. This means that GFs, under their effective conditions, do not conduct the molecular responses of all cells in the same direction, at least in the early stage (5 min) of stimulation. By contrast, effective GFs produce variable responses by single cells. At a later stage, cells that responded appropriately under the specific conditions may be selected and amplified to express the collective cellular response. Alternatively, the stimulation of vigorously changing internal states in single cells with time may be used to search for the appropriate direction of the fate change. These phenomena suggest that a bet-hedging strategy [3] is used for the cell fate decision. Global activation of the multiple elements in the largest component of PCA (PC1; Fig 3) probably indicates the bet-hedging nature of the network. In addition, the response of MCF7 cells in the PC2 direction was opposite for EGF (proliferation) and HRG (differentiation). This difference could be used for the later selection of the cell fate. However, even in this case, overlapping of the single-cell response distributions for EGF and HRG was large especially for B1/B2 and B1/B4 combinations. The distributions were most clearly separated for the B1/B3 combination, supporting the findings of recent studies suggesting that ERBB3 is a key factor in fate determination [24]. Nevertheless, the bet-hedging properties of the cell populations under conditions of a clear cell fate change indicate that exploration of the wide state-space is required for the fate changes.

We previously studied cell fate changes in PC12 cells induced by EGF or NGF [5]. Both of these GFs increased the fate change rates of proliferation and differentiation in single cells. Thus, the specific effects of EGF and NGF—the induction of proliferation and differentiation, respectively—were population behavior. For example, NGF increased the number of differentiated cells by increasing the rate of differentiation by more than the rate of proliferation and by the suppression of dedifferentiation, but not by specifically stimulating cell differentiation or suppressing cell proliferation. We have also used Raman spectral microscopy previously to study the single-cell dynamics of the chemical state in MCF7 cells after stimulation with EGF or HRG [25]. After stimulation, the random temporal fluctuations in the single cells increased, with gradual shifts in the average spectrum for the cell population. These experiments were also suggestive of a bet-hedging strategy in cell fate decisions, regulated by the RAS–MAPK system.

An important question that necessarily arises from our current findings is the nature of the mechanism that produces the variation in cell responses required for bet-hedging. This variation might be the inevitable result of random noise [2] caused by the small numbers of molecules involved in cellular responses [26] and by uncontrollable environmental fluctuations [4]. We evaluated the size of the random noise through a statistical analysis of randomized data. The entropies of the cluster distributions of the single-cell responses increased after randomization in the regions of short and medium threshold distances (Fig. S9B), indicating that the variation was more suppressed in true cells than would be expected for the random activation of molecules. This suppression must therefore be the effect of the correlation (non-zero MI) between molecular activation events. However, under conditions that stimulate clear cell fate changes, the entropies in true cells were larger in regions of long threshold distances than those after randomization. Both the number and probability distribution of the clusters determine the size of the entropy, i.e., the entropy value increases as the number of clusters increases and/or the distribution becomes more even. The large entropies we observed in true cells in the long threshold regions (Fig. S9B) were caused by the longer tails of the cluster number distributions (Fig. 5A, B) i.e., the wider dynamic ranges in the multidimensional space of the molecular responses, which were observed under the conditions that induce cell fate changes (Fig. 5C).

In conclusion, the cell-to-cell response variation in true cells is not simply determined by random noise, but is structured by the signal transduction network to reduce the randomness and increase the dynamic range of the response. In other words, cell-to-cell diversity is designed by the signal transduction network. This increased single-cell diversity seems to be generated by the use of various signal transduction pathways in bush-like networks.

## Materials and Methods

### Preparation of cells

HeLa and A431 cells were obtained from RIKEN Cell Bank and MCF7 cells from the American Type Culture Collection. All cells were maintained in Dulbecco’s modified Eagle’s medium supplemented with 10% fetal bovine serum (FBS) at 37°C under 5% CO2. Cells were cultured for 1 day after transfer and then starved of FBS for 18 h before use in the experiments.

### Antibodies

The primary antibodies used were: rabbit anti-pERBB1 IgG (53A5, #4407, CST), rabbit anti-pERBB2 IgG (6B12, #2243, CST), rabbit anti-pERBB3 IgG (21D3, #4791, CST), rabbit anti-pERBB4 IgG (21A9, #4757, CST), mouse anti-pCaMKII IgG (22B1, MA1-047, Thermo Fisher), mouse anti-PIP2 IgM (2C11, sc-53412, Santa Cruz Biotechnology), and mouse anti-ppERK IgG1 (E10, #9106, CST).

The secondary antibodies used for western blotting were: horseradish peroxidase (HRP)-conjugated anti-rabbit IgG (#7074, CST) and anti-mouse IgG (#7076, CST).

The fluorophore-conjugated secondary antibodies used were: Atto-425-conjugated goat anti-mouse IgG (610-151-121S, Rockland Antibodies), tetramethylrhodamine (TRITC)-conjugated anti-mouse IgM (#731781, Beckman Coulter), and Alexa-594-conjugated anti-rabbit IgG (A11012, Molecular Probes).

For multicolor immunofluorescent staining, the anti-pERBB1 and anti-ppERK antibodies were directly conjugated with Alexa 488 and Alexa 647, respectively, with the Microscale Protein Labeling Kit (Molecular Probes), in accordance with the manufacturer’s instructions.

### Western blotting analysis of protein phosphorylation

Cells at 70% confluence after FBS starvation were incubated with 0–30 nM (final concentration) murine EGF (PeproTech) or recombinant human NRG-β1/HRG-β1 EGF domain (HRG; R&D Systems) for 5 min at 25°C, washed, and harvested in SDS sampling buffer containing 1 mM Na3VO4. The proteins in the lysates were separated by SDS-PAGE and transferred to polyvinylidene difluoride membranes (BD Biosciences). The phosphorylation of the ERBBs and ERK was detected on the membranes with anti-pERBB1–B4 and anti-ppERK primary antibodies, respectively, and an HRP-linked secondary antibody. Antibody binding was detected using the ECL Prime Western Blotting Detection Reagent (Amersham), and chemiluminescence was measured with a luminescent image analyzer (LAS 500, GE Healthcare).

### Immunofluorescence staining

For immunofluorescence straining, each cell type was plated on a 9 × 9 mm^2^ glass coverslip in a well of a 24-well cell culture plate, at a density of 10^4^ cells/well. After culture for 1 day in 10% serum followed by serum starvation for 18 h, the cells were stimulated with 20 nM EGF or 30 nM HRG for 5 min at 25 °C. They were then fixed, permeabilized, and stained simultaneously with anti-pERBB1, and one of anti-pErbB2– B4, anti-pCaMKII, anti-PIP2, or anti-ppERK antibodies. Either the direct or indirect immunofluorescence method with the appropriate secondary antibody was used (see Fig. S2 for details). The procedures after cell stimulation were performed with a microplate dispenser (MultiFlo FX, BioTek). After immunostaining, the coverslips were mounted in Abberior Mount Solid (Abberior).

### Fluorescence microscopy and spectral unmixing

Micrographs of the cells were acquired under a fluorescence microscope (IX81, Olympus) equipped with a 20× objective (UPLSAPO, Olympus) and a 130 W Hg lamp (U-HGLGPS, Olympus), with a cMOS camera (ORCA-Flash4.0, Hamamatsu). By changing the filter cubes, six images were obtained successively with different excitation and emission channels (five for fluorescent dyes and one for AF) in each field of view (filters are listed in Table S1).

After shading correction and background subtraction, the fluorescence signals in the six channels were unmixed into five single-dye signals and AF, based on the fluorescence spectrum of each dye and AF, measured under a microscope. Spectral unmixing was performed by multiplying the inverse of the 6 × 6 fluorescence spectra matrix by each pixel in the stack of six channels. Isolated cells were selected randomly, and the average signal intensity per unit area in each single cell was measured and used in subsequent analyses. Image calculations and measurements were made using ImageJ software (NIH).

### Multi-component and statistical analyses

The molecular responses in single cells were calculated as the difference between the immunofluorescence intensities in each cell with GF stimulation and the average intensity in the cells without stimulation. The difference values were normalized with their average and SD (Mahalanobis’ distance) and used for the subsequent analyses. PCA was performed using the prcomp function of R (https://www.R-project.org/) with maximization of the variance.

The definition of mutual information (MI) is as follows:

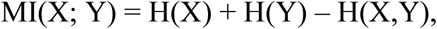

where H(X) is the information entropy of the probability distribution, P(x), of X:

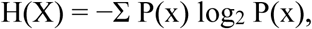

and H(X, Y) is the joint entropy of P(x) and P(y):

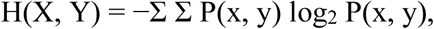

[27]. P(x, y) is the joint probability of X and Y. The values of MI were calculated with the mutinformation function [28] of R with the empirical probability distribution estimator. Prior to the calculation, the distributions of the single-cell responses were discretized with a bin size of 1/2 SD of each distribution. This bin size (1/2 SD) was chosen empirically but was larger than the noise during signal detection (Fig. S6), and the difference in the amounts of MI for the true and randomized data was maximal at a bin size of around 1/2 SD (Fig. S7).

Hierarchical clustering of the five-channel staining patterns of single cells was performed with Ward’s method, using the ward.D2 method in the hclust function of R. The normalized response intensities from three independent experiments were gathered and used for clustering. Shannon’s entropy (Fig. S9) is the information entropy of the cluster distribution, for which the logarithm at base 10 was chosen.

## Supporting information

Miyagi et al supplement

## Acknowledgments

We thank Atsushi Mochizuki at Kyoto University and Shinsuke Uda at Kyushu University for critical discussions.

## Author contributions

Y.S. designed the research; H.M performed the experiments; H.M. and Y.S. analyzed the data; all authors wrote the manuscript.

