## Supplementary material for "Cell-to-cell diversification in ERBB-RAS-MAPK signal transduction that produces cell-type specific growth factor responses": Miyagi et al supplement

**Supplementary Figures and Tables**

**Figure S1**. Dose responses of ERBBs and ERK to growth factor stimulations

HeLa, A431, and MCF7 cells were stimulated with various concentrations of EGF or HRG for 5 min at 25°C. The phosphorylation of the direct receptors of each growth factor or ERK was analyzed by western blotting. Because the expression levels were low, dose–response measurements were not made for ERBB3 in HeLa cells, ERBB4 in A431 cells, or ERBB1 in MCF7 cells. Instead, EGF stimulation of MCF7 cells was measured as ERK phosphorylation. (A) Immunostaining results. Numbers indicate the concentrations of EGF or HRG (nM). The upper and lower arrowheads indicate 225 and 150 kDa for ERBBs respectively, and 52 and 38 kDa for ERK, respectively. (B) Dose–response curves. Staining intensities were normalized to the maximum value. Results are fitted with the Hill function to obtain the indicated $EC_{50}$ values.

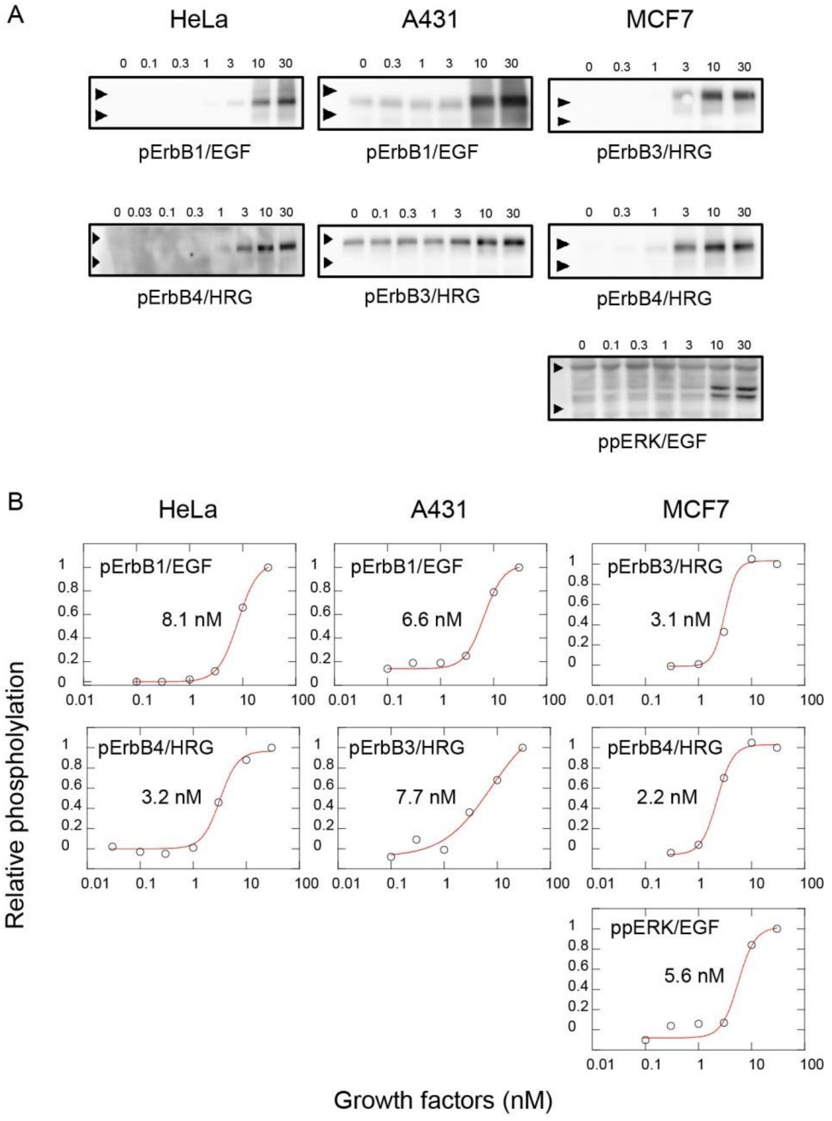

**Figure S2.** Protocol for five-color immunofluorescent staining

All incubations were at 25°C unless otherwise stated. Hank's balanced salt solution without glucose was used as the solvent for the reagents and as a washing solution. Antibodies (40 μl/coverslip) were applied by hand. A microplate dispenser was used for all other procedures. Further details of this methodology can be accessed online at dx.doi.org/10.17504/protocols.io.9gxh3xn

PFA: paraformaldehyde.

Cells on a 9 × 9 mm coverslip after growth factor simulation (in a single well of a 24-well plate)

↓

**Fixation 1:** 400 μl, 3% PFA 1 mM $Na_3VO_4$ 1 mM NaF, 10 min

↓ Wash

**Permeablization 1:** 400 μl, 50 μg/ml digitonin in 3% PFA 1 mM $Na_3VO_4$ 1 mM NaF, 10 min

↓ Wash

**Blocking 1:** 300 μl, Blocking One-P (Nacalai), 30 min

↓

**1º Ab 1:** 1/100 anti-pCaMKII in Blocking One-P, 60 min

↓ Wash

**2º Ab 1:** 1/600 ATTO425-anti-mouse IgG in Blocking One-P, 30 min

↓ Wash

**1º Ab 2:** 1/600 anti-$PIP_2$ in Blocking One-P, 60 min

↓ Wash

**Fixation 2:** 400 μl, 3% PFA, 10 min

↓ Wash

**Permeablization 2:** 400 μl 0.01% Triton X-100 in 3% PFA, 1 mM $Na_3VO_4$ 1 mM NaF, 10 min

↓ Wash

**Blocking 2:** 300 μl, Blocking One-P, 30 min

↓

**1º Ab 3:** 1/600 anti-pERBB2~4 in Blocking One-P, overnight at 4 ºC

↓ Wash

**2º Ab 2:** 1/3600 Alexa-594-conjugated anti-rabbit IgG + 1/2400 TRITC-anti-mouse IgM in Blocking One-P, 30 min

↓ Wash

**1º Ab 4:** 1/300 Alexa-488-conjugated anti-pERBB1 + 1/300 Alexa594-anti-ppERK in Blocking One-P, 60 min

↓ Wash

**Mount:** 4 μl Abberior Mount Solid (Abberior instruments), overnight at 4 ºC

**Figure S3**. Spectral unmixing

(A) Emission spectra of the dye solutions in mounting medium and the autofluorescence (AF) of cells were measured under a fluorescence microscope. Dye and AF spectra were normalized to the peak and AF channel, respectively. (B) Typical six-channel images of cells stained with single dyes before (upper) and after (lower) spectral unmixing. (C) The signal intensity in each channel before (blue) and after (red) spectral unmixing is shown as the average ± SD of single-cell measurements. After unmixing, significant signals remained only in the AF and corresponding dye channels. SD in the corresponding dye channel indicates cell-to-cell deviation in the molecular response. SD in the other channels was reduced after unmixing. TRITC and Alexa 647 were examined in HeLa and MCF7 cells, respectively. Other dyes were examined in A431 cells. Cells (n = 17–20) were measured under each condition. AU, arbitrary unit.

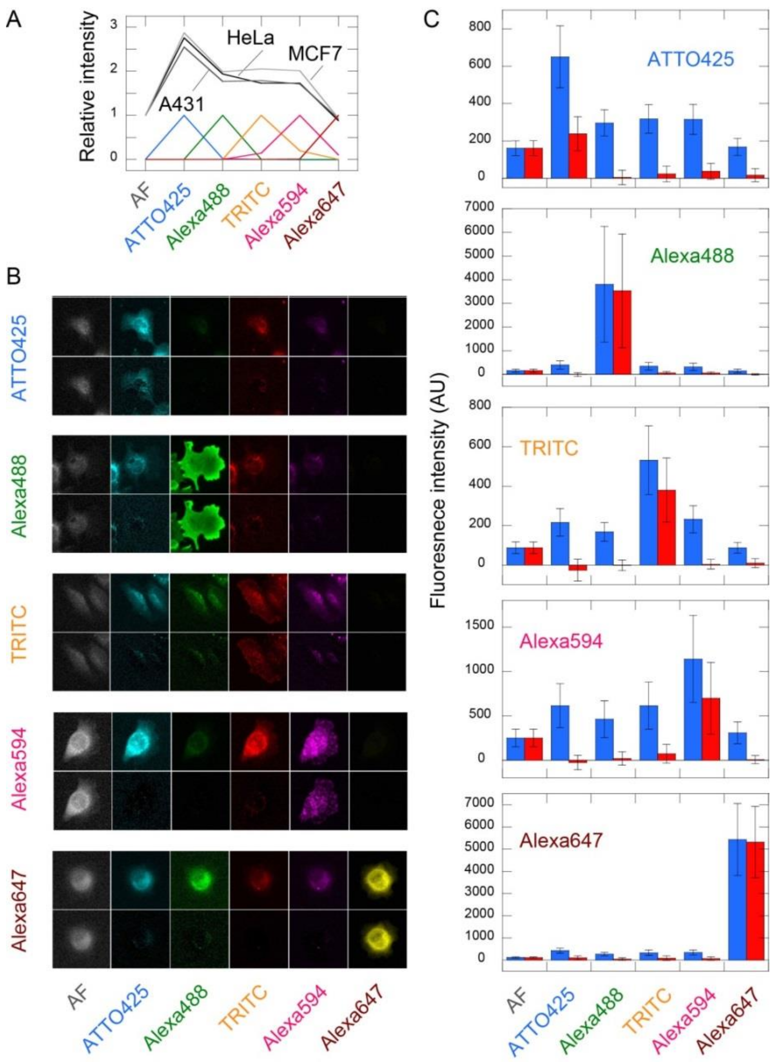

**Figure S4.** Response to growth factor stimulation in the averaged cells

(A) Typical five-color staining micrographs after unmixing. The upper left images of the HeLa cells are the same as those shown in Figure 1A. (B–D) Fold changes in the staining intensity are shown as the average (± SD) for 115–135 randomly selected cells. Three independent experiments were performed under the same conditions. Significance levels against cells without GF stimulation are indicated: *P < 0.05, **P < 0.01 (*t* test).

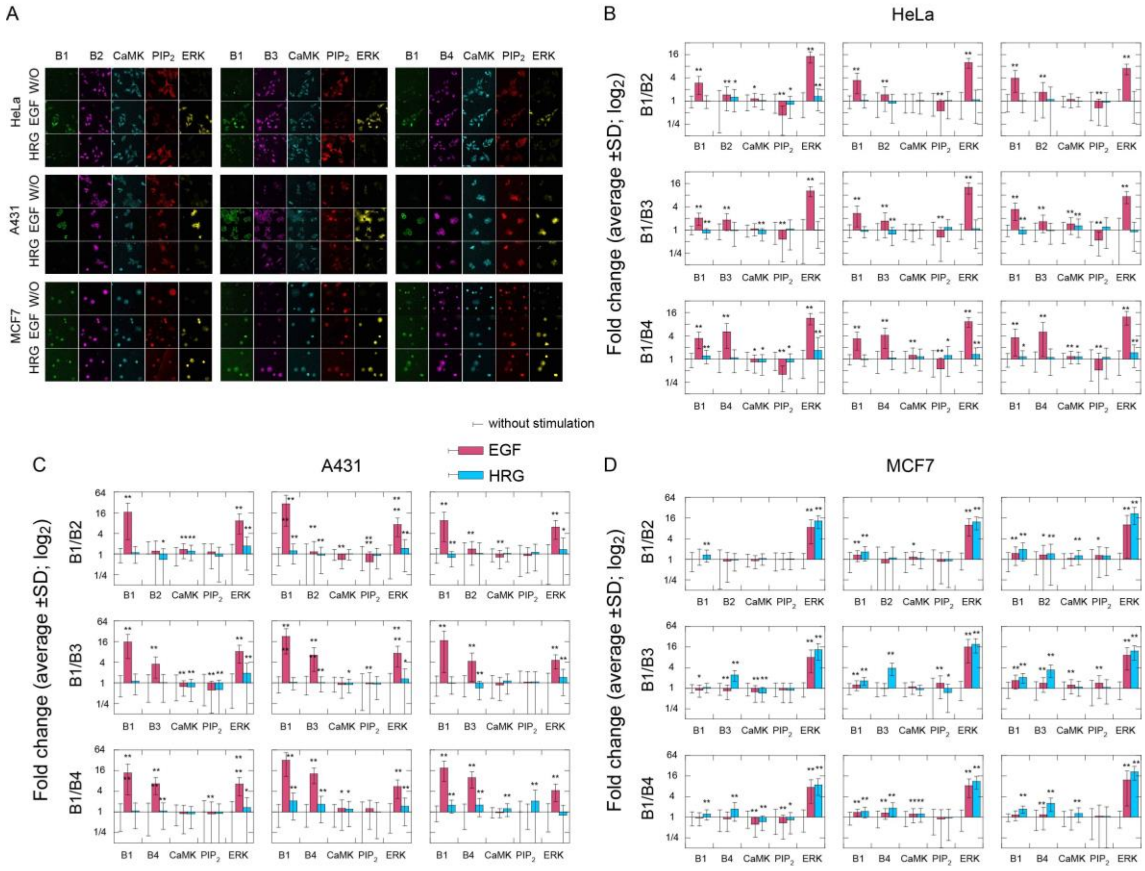

**Figure S5.** Single-cell distributions of molecular responses

Results of all experiments shown in Figure 2 for HeLa (A), A431 (B), and MCF7 (C) cells. Intensities were normalized with the average of all cells with and without stimulation. Diagonal panels show the intensity distributions in one dimension. Vertical axes in the diagonal panels indicate the numbers of cells. Data are shown for cells without stimulation (green) or with EGF (red) or HRG (blue) stimulation.

Fig. S5A HeLa

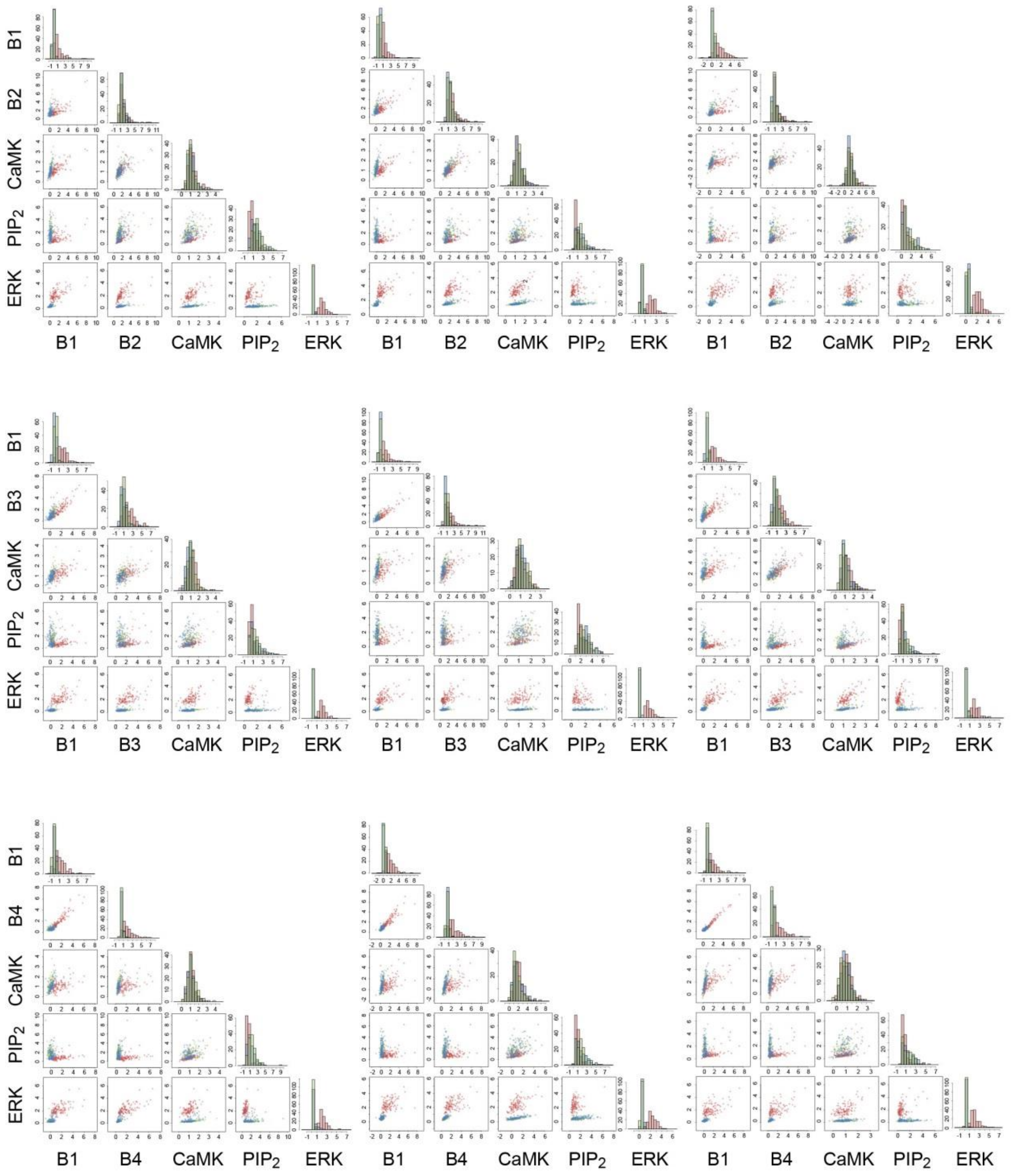

Fig. S5B    A431

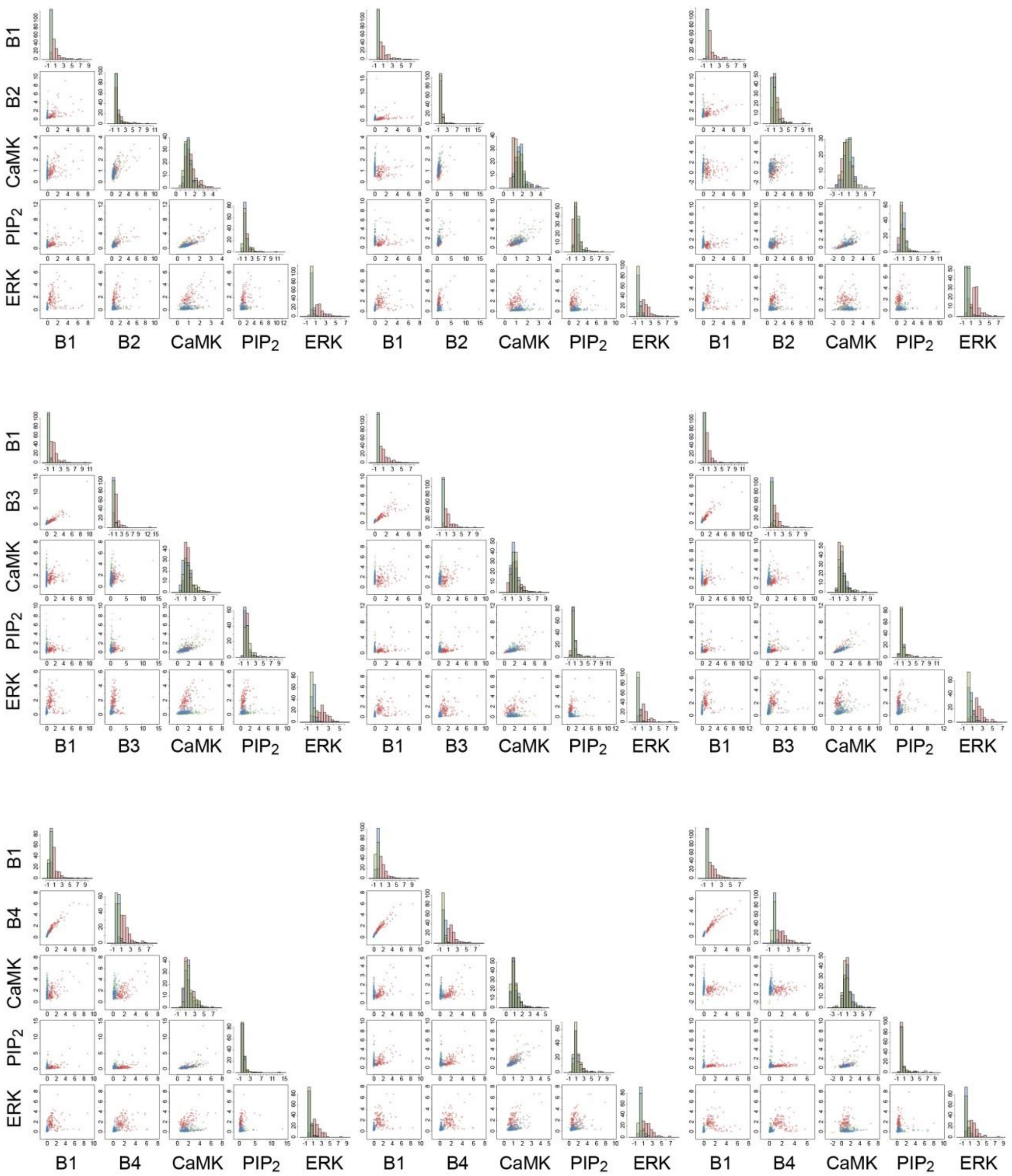

Fig. S5C MCF7

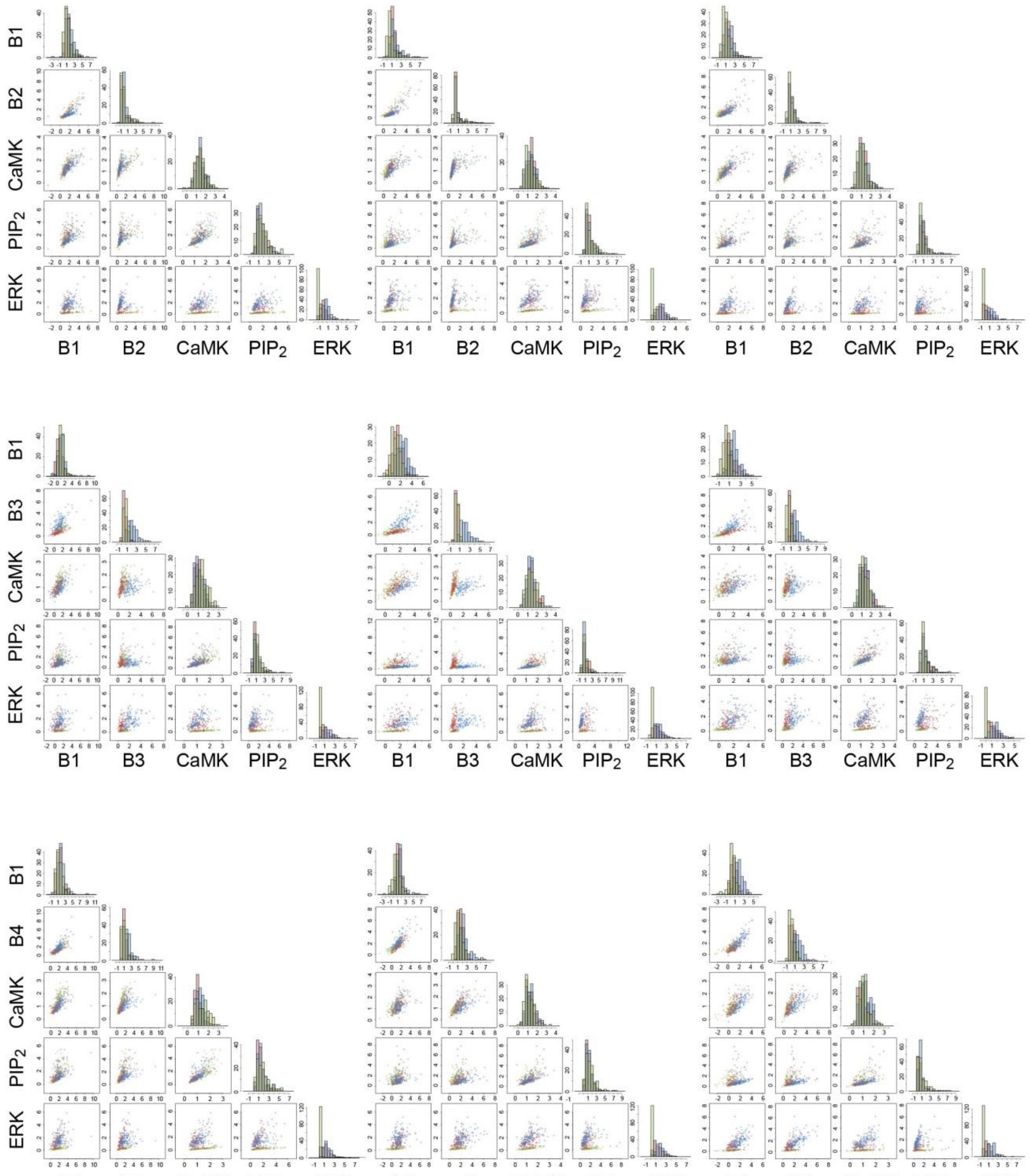

**Figure S6.** Measurement errors in the immunofluorescence detection experiments

Molecules in the same single cells were detected with a mixture of the same antibodies conjugated with different fluorescent dyes. Correlations between the two colors are shown in the upper panels, where the staining intensities are plotted as the differences from the average value normalized using the SD. A correlation coefficient (R) of nearly 1 indicates that there was little noise during the staining and detection. In the lower panels, measurement errors (dx) were plotted against the estimated true signal intensities (x). Definitions of x and dx are shown in the panel for the pERBB2 distribution. We assumed that the noise was evenly shared by the two channels. Numbers in the lower panels indicate the SD of the dx (measurement error in the unit SD of the signal intensities). Data was obtained in HeLa (ERBB1, ERBB4, $PIP_2$), A431 (ERBB3, ERK), and MCF7 (ERBB2, CaMKII) cells stimulated with EGFs.

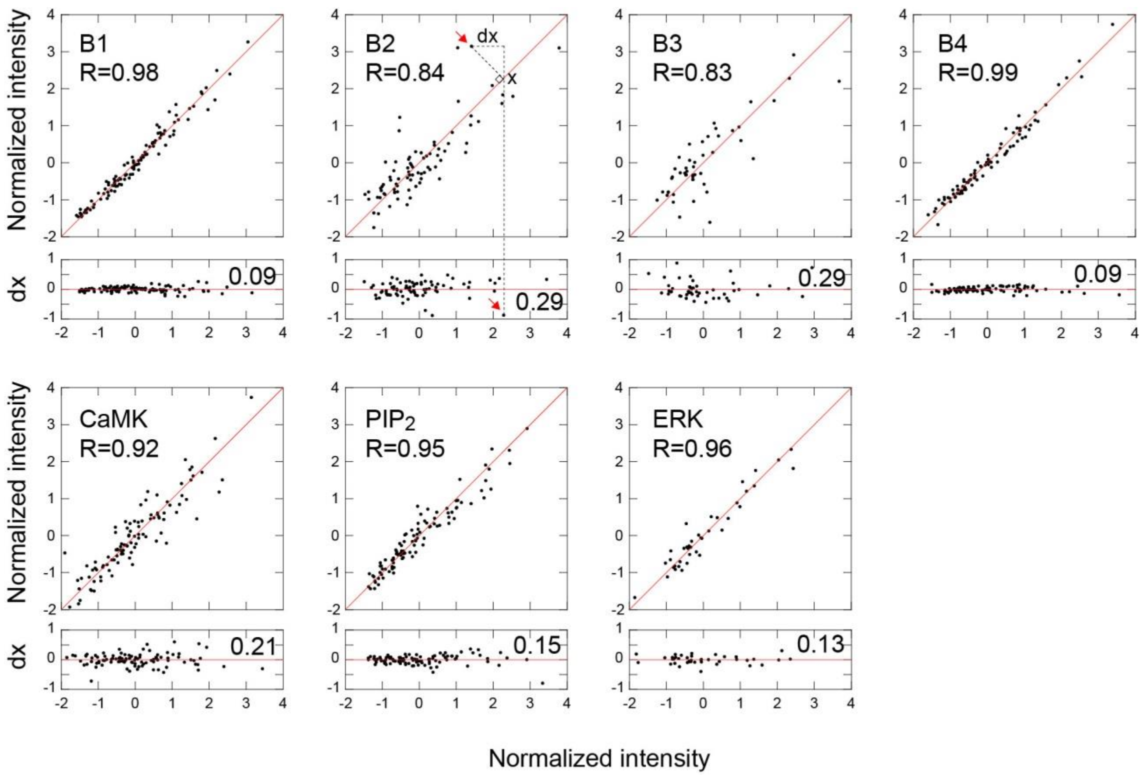

**Figure S7.** Bin-size dependence of the mutual information (MI) amounts

The MI amounts for the true (upper panels) and randomized (middle panels) data were plotted against the bin size to discretize the probability distributions. Randomization was performed by permuting the response intensities of each element between single cells. The unit of the bin size is the SD in each distribution. Differences in MI amounts (the upper panels minus the middle panels) are shown in the lower panels. Each point shows the average value from the independent experiments. The differences were maximal at around a bin size of 1/2 SD, in general, for combinations of molecules and the conditions of the experiments. Red and blue spots indicate results for EGF and HRG stimulations, respectively. MI values were positive (non-zero) even after randomization because the number of data points was limited.

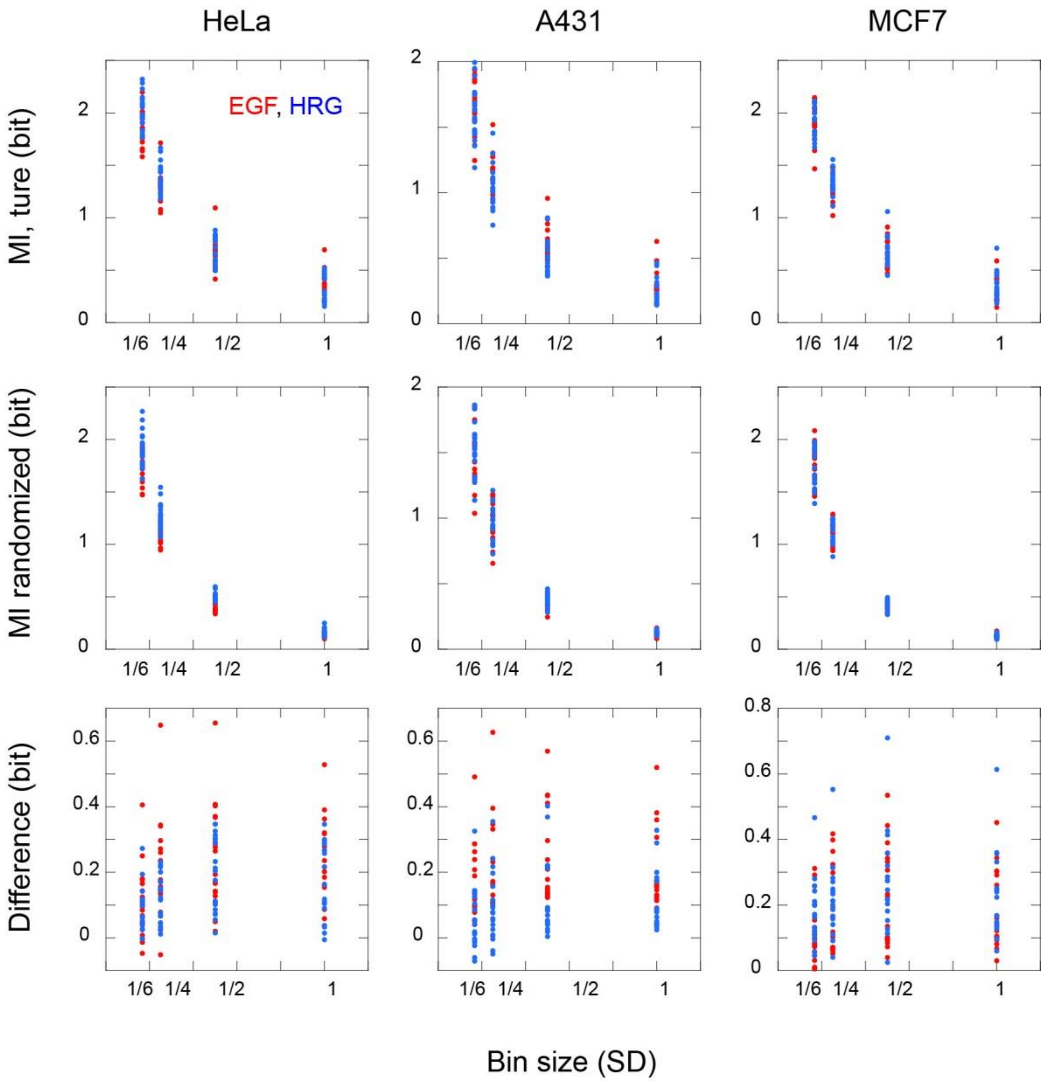

**Figure S8.** Response intensity and MI

Response intensities, defined as the vector length of the average of two elements, $(X_{ave}^2 + Y_{ave}^2)^{1/2}$, were compared to the MI between the same two elements. Here, $X_{ave}$ and $Y_{ave}$ are the average fold changes in the responses of X and Y elements to GFs, respectively, as shown in Figures 1 and S4. The averages of three independent experiments are shown ± SE.

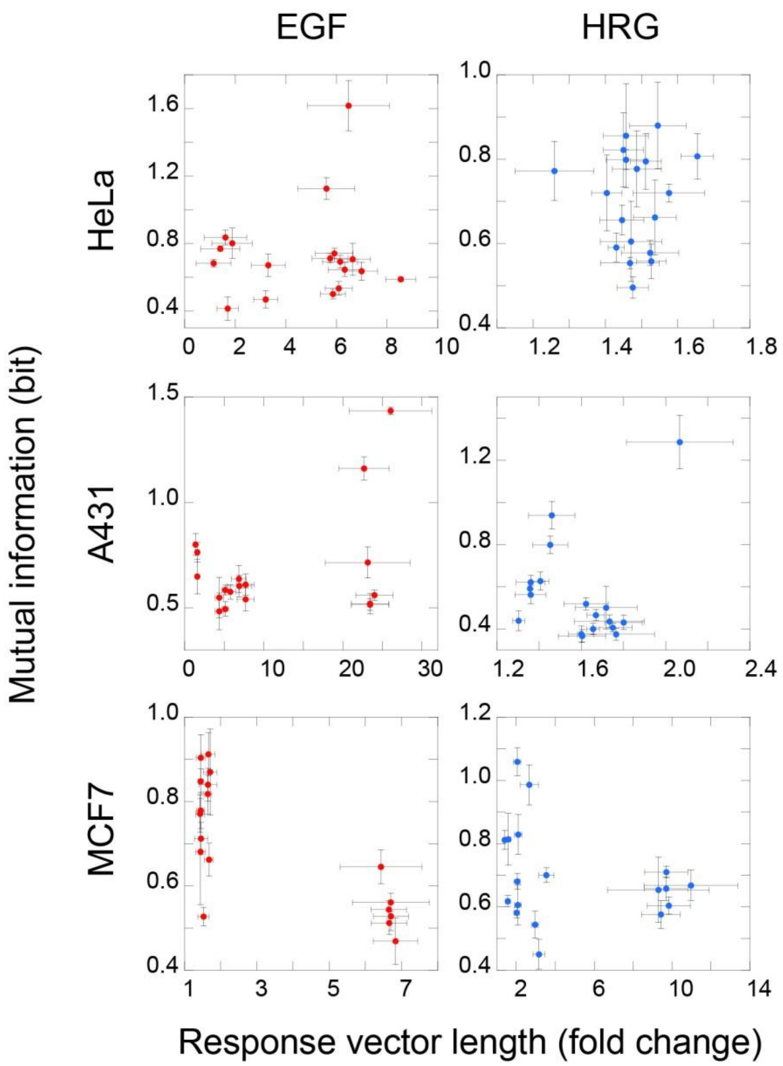

**Figure S9.** Shannon's entropy was calculated for the cluster distributions as a function of the threshold distance. Results were compared between the true datasets for EGF and HRG (A), or between the true and randomized datasets (B). AU: arbitrary unit.

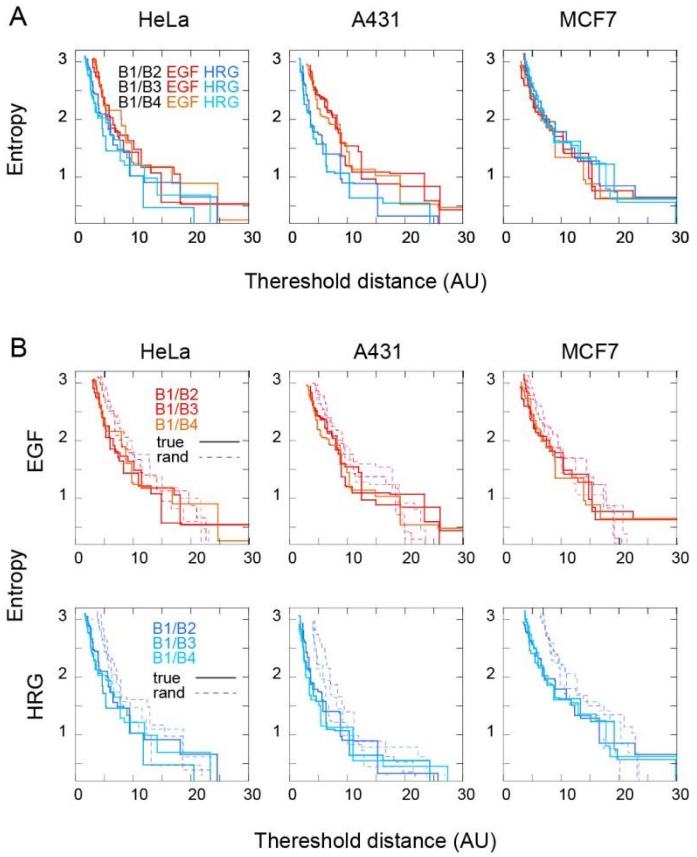

**Table S1. Filter sets used for multicolor fluorescence imaging**

| Dye (channel) | Filter set | Ex | DM | Em |
|---|---|---|---|---|
| Autofluorescence | Olympus | BP330-385 | 405 | 440DF20 |
| Atto 425 | Chroma 49001 | 436/20 | 455LP | 480/40m |
| Alexa 488 | Olympus U-MNIBA2 | BP470–490 | 505 | 510–550 |
| TRITC | Olympus U-MRFPHQ | BP535–555 | 565 | 570–625 |
| Alexa 594 | Chroma49306 | 580/25λ | T6001pxr | ET625/30m |
| Alexa 647 | Chroma U-M4100BHQ | 620/60λ | 660LP | 700/75m |

Ex, excitation filter; DM, dichroic mirror; Em, emission filter

**Table S2. Mutual information amounts**

Average and SE values for mutual information (bits) from three or nine independent measurements. P values were calculated using a *t* test against randomized data.

| HeLa | | EGF | | | | HRG | | | |
|---|---|---|---|---|---|---|---|---|---|
| Molecule1 | Molecule2 | Average | SE | n | p | Average | SE | n | p |
| B1 | B2 | 0.707 | 0.094 | 3 | 0.020 | 0.662 | 0.089 | 3 | 0.050 |
| B1 | B3 | 1.126 | 0.065 | 3 | 0.001 | 0.772 | 0.070 | 3 | >0.1 |
| B1 | B4 | 1.618 | 0.148 | 3 | 0.001 | 0.720 | 0.021 | 3 | 0.001 |
| B1 | CaMKII | 0.742 | 0.030 | 9 | 0.001 | 0.799 | 0.024 | 9 | 0.002 |
| B2 | CaMKII | 0.837 | 0.044 | 3 | 0.002 | 0.822 | 0.088 | 3 | 0.050 |
| B3 | CaMKII | 0.802 | 0.090 | 3 | 0.020 | 0.880 | 0.102 | 3 | 0.050 |
| B4 | CaMKII | 0.672 | 0.066 | 3 | 0.050 | 0.856 | 0.123 | 3 | 0.050 |
| B1 | PIP2 | 0.501 | 0.028 | 9 | 0.020 | 0.496 | 0.025 | 9 | >0.1 |
| B2 | PIP2 | 0.770 | 0.016 | 3 | 0.002 | 0.591 | 0.034 | 3 | >0.1 |
| B3 | PIP2 | 0.415 | 0.069 | 3 | >0.1 | 0.605 | 0.095 | 3 | >0.1 |
| B4 | PIP2 | 0.469 | 0.051 | 3 | >0.1 | 0.554 | 0.014 | 3 | 0.100 |
| B1 | ERK | 0.589 | 0.013 | 9 | 0.050 | 0.578 | 0.029 | 9 | 0.050 |
| B2 | ERK | 0.713 | 0.028 | 3 | 0.005 | 0.777 | 0.090 | 3 | 0.050 |
| B3 | ERK | 0.646 | 0.040 | 3 | 0.005 | 0.720 | 0.090 | 3 | >0.1 |
| B4 | ERK | 0.636 | 0.054 | 3 | 0.050 | 0.807 | 0.054 | 3 | 0.005 |
| CaMKII | PIP2 | 0.683 | 0.022 | 9 | 0.001 | 0.656 | 0.035 | 9 | 0.020 |
| CaMKII | ERK | 0.693 | 0.037 | 9 | 0.020 | 0.795 | 0.066 | 9 | 0.010 |
| PIP2 | ERK | 0.535 | 0.039 | 9 | 0.100 | 0.558 | 0.041 | 9 | >0.1 |

| A431 | | EGF | | | | HRG | | | |
|---|---|---|---|---|---|---|---|---|---|
| Molecule1 | Molecule2 | Average | SE | n | p | Average | SE | n | p |
| B1 | B2 | 0.716 | 0.073 | 3 | 0.005 | 0.622 | 0.033 | 3 | 0.020 |
| B1 | B3 | 1.162 | 0.054 | 3 | 0.001 | 0.939 | 0.065 | 3 | 0.005 |
| B1 | B4 | 1.435 | 0.018 | 3 | 0.001 | 1.287 | 0.127 | 3 | 0.005 |
| B1 | CaMKII | 0.516 | 0.028 | 9 | 0.001 | 0.519 | 0.029 | 9 | 0.002 |
| B2 | CaMKII | 0.649 | 0.082 | 3 | 0.005 | 0.628 | 0.043 | 3 | 0.020 |
| B3 | CaMKII | 0.549 | 0.095 | 3 | >0.1 | 0.592 | 0.029 | 3 | 0.002 |
| B4 | CaMKII | 0.611 | 0.051 | 3 | 0.050 | 0.436 | 0.031 | 3 | >0.1 |
| B1 | PIP2 | 0.520 | 0.048 | 9 | 0.020 | 0.367 | 0.030 | 9 | >0.1 |
| B2 | PIP2 | 0.764 | 0.047 | 3 | 0.005 | 0.563 | 0.044 | 3 | 0.005 |
| B3 | PIP2 | 0.484 | 0.088 | 3 | 0.050 | 0.439 | 0.046 | 3 | >0.1 |
| B4 | PIP2 | 0.541 | 0.055 | 3 | >0.1 | 0.375 | 0.028 | 3 | >0.1 |
| B1 | ERK | 0.561 | 0.024 | 9 | 0.050 | 0.431 | 0.034 | 9 | 0.050 |
| B2 | ERK | 0.577 | 0.035 | 3 | 0.020 | 0.376 | 0.038 | 3 | >0.1 |
| B3 | ERK | 0.638 | 0.064 | 3 | 0.050 | 0.501 | 0.102 | 3 | >0.1 |
| B4 | ERK | 0.604 | 0.051 | 3 | 0.020 | 0.407 | 0.020 | 3 | >0.1 |
| CaMKII | PIP2 | 0.801 | 0.052 | 9 | 0.001 | 0.799 | 0.042 | 9 | 0.020 |
| CaMKII | ERK | 0.585 | 0.023 | 9 | 0.020 | 0.466 | 0.027 | 9 | 0.050 |
| PIP2 | ERK | 0.495 | 0.034 | 9 | 0.100 | 0.400 | 0.027 | 9 | 0.050 |

| MCF7 | | EGF | | | | HRG | | | |
|---|---|---|---|---|---|---|---|---|---|
| Molecule1 | Molecule2 | Average | SE | n | p | Average | SE | n | p |
| B1 | B2 | 0.912 | 0.051 | 3 | 0.001 | 1.059 | 0.044 | 3 | 0.001 |
| B1 | B3 | 0.840 | 0.039 | 3 | 0.001 | 0.701 | 0.023 | 3 | 0.001 |
| B1 | B4 | 0.870 | 0.102 | 3 | 0.010 | 0.986 | 0.063 | 3 | 0.002 |
| B1 | CaMKII | 0.818 | 0.047 | 9 | 0.001 | 0.829 | 0.063 | 9 | 0.002 |
| B2 | CaMKII | 0.779 | 0.043 | 3 | 0.001 | 0.814 | 0.082 | 3 | 0.005 |
| B3 | CaMKII | 0.771 | 0.044 | 3 | 0.010 | 0.544 | 0.043 | 3 | 0.100 |
| B4 | CaMKII | 0.904 | 0.055 | 3 | 0.002 | 0.681 | 0.025 | 3 | 0.005 |
| B1 | PIP2 | 0.662 | 0.039 | 9 | 0.020 | 0.606 | 0.063 | 9 | >0.1 |
| B2 | PIP2 | 0.712 | 0.038 | 3 | 0.005 | 0.618 | 0.018 | 3 | 0.001 |
| B3 | PIP2 | 0.527 | 0.022 | 3 | 0.100 | 0.450 | 0.047 | 3 | >0.1 |
| B4 | PIP2 | 0.681 | 0.126 | 3 | 0.100 | 0.582 | 0.017 | 3 | 0.050 |
| B1 | ERK | 0.528 | 0.034 | 9 | >0.1 | 0.604 | 0.027 | 9 | 0.050 |
| B2 | ERK | 0.469 | 0.055 | 3 | >0.1 | 0.668 | 0.048 | 3 | 0.005 |
| B3 | ERK | 0.561 | 0.021 | 3 | 0.050 | 0.576 | 0.044 | 3 | 0.050 |
| B4 | ERK | 0.645 | 0.040 | 3 | 0.005 | 0.654 | 0.103 | 3 | >0.1 |
| CaMKII | PIP2 | 0.848 | 0.029 | 9 | 0.001 | 0.812 | 0.030 | 9 | 0.020 |
| CaMKII | ERK | 0.544 | 0.021 | 9 | >0.1 | 0.710 | 0.019 | 9 | 0.010 |
| PIP2 | ERK | 0.512 | 0.027 | 9 | 0.050 | 0.658 | 0.037 | 9 | 0.010 |

**Table S3. Single-cell response patterns**

The average (Ave) and SD values for the five-molecule responses of each cluster shown in Figure 4D are listed. n, number of cells.

A. ERBB1/B2

HeLa + EGF

| | ERBB1 | | ERBB2 | | CaMKII | | PIP2 | | ERK | | | |
|---|---|---|---|---|---|---|---|---|---|---|---|---|
| Cluster | Ave | SD | Ave | SD | Ave | SD | Ave | SD | Ave | SD | n | % |
| 1 | -0.30 | 0.29 | -0.66 | 0.26 | -0.92 | 0.39 | -0.78 | 0.27 | 0.29 | 0.39 | 102 | 28.0 |
| 2 | 0.21 | 0.40 | -0.14 | 0.42 | -0.21 | 0.43 | -0.60 | 0.25 | 0.70 | 0.51 | 118 | 32.4 |
| 3 | -0.06 | 0.36 | 0.01 | 0.43 | -0.34 | 0.55 | 0.54 | 0.48 | 0.70 | 0.43 | 31 | 8.5 |
| 4 | 1.60 | 0.58 | -0.07 | 0.20 | 0.23 | 0.54 | -0.57 | 0.37 | 0.80 | 0.42 | 30 | 8.2 |
| 5 | 1.13 | 0.73 | 1.31 | 0.74 | 0.76 | 0.64 | 0.31 | 0.70 | 1.16 | 0.65 | 55 | 15.1 |
| 6 | 2.51 | 1.07 | 1.85 | 1.13 | 2.20 | 0.90 | 0.66 | 1.05 | 1.89 | 0.71 | 25 | 6.9 |
| 7 | 6.11 | 0.22 | 6.43 | 0.61 | 3.66 | 0.54 | 2.62 | 0.56 | 2.81 | 0.65 | 3 | 0.8 |

HeLa + HRG

| | ERBB1 | | ERBB2 | | CaMKII | | PIP2 | | ERK | | | |
|---|---|---|---|---|---|---|---|---|---|---|---|---|
| Cluster | Ave | SD | Ave | SD | Ave | SD | Ave | SD | Ave | SD | n | % |
| 1 | -0.67 | 0.15 | -0.64 | 0.26 | -0.41 | 0.51 | -0.54 | 0.28 | -1.02 | 0.15 | 105 | 35.7 |
| 2 | -0.69 | 0.15 | -0.32 | 0.55 | -0.47 | 0.40 | 0.39 | 0.47 | -1.01 | 0.21 | 80 | 27.2 |
| 3 | -0.68 | 0.14 | -0.03 | 0.53 | 0.16 | 0.66 | 2.25 | 0.89 | -0.92 | 0.17 | 51 | 17.3 |
| 4 | -0.42 | 0.28 | 0.34 | 0.71 | 1.06 | 0.42 | -0.01 | 0.47 | -0.81 | 0.21 | 42 | 14.3 |
| 5 | -0.33 | 0.20 | 1.16 | 0.76 | 2.59 | 0.84 | 0.90 | 0.54 | -0.55 | 0.40 | 13 | 4.4 |
| 6 | -0.17 | 0.19 | 4.27 | 1.92 | 2.70 | 0.29 | 3.05 | 0.46 | -0.63 | 0.07 | 3 | 1.0 |

A431 + EGF

| | ERBB1 | | ERBB2 | | CaMKII | | PIP2 | | ERK | | | |
|---|---|---|---|---|---|---|---|---|---|---|---|---|
| Cluster | Ave | SD | Ave | SD | Ave | SD | Ave | SD | Ave | SD | n | % |
| 1 | -0.09 | 0.21 | -0.53 | 0.20 | -0.97 | 0.35 | -0.74 | 0.24 | 0.29 | 0.40 | 69 | 21.0 |
| 2 | 0.18 | 0.28 | 0.17 | 0.42 | -0.71 | 0.41 | -0.51 | 0.28 | 1.16 | 0.42 | 40 | 12.2 |
| 3 | 0.33 | 0.54 | -0.19 | 0.41 | -0.29 | 0.52 | -0.07 | 0.33 | -0.12 | 0.44 | 38 | 11.6 |
| 4 | 0.12 | 0.33 | -0.20 | 0.38 | 0.36 | 0.57 | -0.19 | 0.31 | 0.89 | 0.45 | 77 | 23.5 |
| 5 | 0.48 | 0.38 | 1.08 | 0.75 | 0.35 | 0.71 | 1.26 | 0.42 | 0.45 | 0.56 | 11 | 3.4 |
| 6 | 0.48 | 0.46 | 0.59 | 0.49 | 0.75 | 1.10 | 0.39 | 0.54 | 2.76 | 0.76 | 20 | 6.1 |
| 7 | 0.99 | 0.55 | 1.10 | 0.84 | 1.99 | 1.25 | 3.56 | 1.20 | 1.35 | 1.12 | 6 | 1.8 |
| 8 | 0.84 | 0.47 | 3.27 | 1.19 | 0.69 | 1.26 | 1.28 | 0.73 | 1.97 | 1.06 | 11 | 3.4 |
| 9 | 2.09 | 0.71 | 0.20 | 0.55 | -0.51 | 0.71 | -0.32 | 0.48 | -0.47 | 0.39 | 40 | 12.2 |
| 10 | 4.56 | 0.83 | 1.32 | 0.98 | 1.23 | 0.99 | 0.87 | 1.04 | 0.08 | 0.96 | 12 | 3.7 |
| 11 | 2.34 | 0.93 | 6.34 | 1.78 | 3.54 | 1.25 | 4.60 | 2.96 | 3.70 | 1.35 | 4 | 1.2 |

A431 + HRG

| | ERBB1 | | ERBB2 | | CaMKII | | PIP2 | | ERK | | | |
|---|---|---|---|---|---|---|---|---|---|---|---|---|
| Cluster | Ave | SD | Ave | SD | Ave | SD | Ave | SD | Ave | SD | n | % |
| 1 | -0.61 | 0.03 | -0.62 | 0.23 | -0.92 | 0.28 | -0.63 | 0.23 | -0.71 | 0.25 | 76 | 22.8 |
| 2 | -0.60 | 0.04 | -0.35 | 0.40 | -0.23 | 0.32 | -0.12 | 0.40 | -0.68 | 0.27 | 129 | 38.6 |
| 3 | -0.60 | 0.05 | -0.17 | 0.43 | 0.73 | 0.42 | 0.16 | 0.45 | -0.62 | 0.28 | 95 | 28.4 |
| 4 | -0.55 | 0.06 | 0.83 | 0.80 | 1.95 | 0.67 | 0.74 | 0.57 | -0.48 | 0.35 | 17 | 5.1 |
| 5 | -0.59 | 0.03 | 0.97 | 0.93 | 1.71 | 1.07 | 3.09 | 0.98 | -0.44 | 0.72 | 12 | 3.6 |
| 6 | -0.57 | 0.04 | 4.15 | 1.30 | 1.38 | 1.71 | 1.21 | 0.85 | -0.77 | 0.06 | 4 | 1.2 |
| 7 | -0.56 | NA | 3.63 | NA | 2.97 | NA | 9.20 | NA | -0.72 | NA | 1 | 0.3 |

MCF7 + EGF

| | ERBB1 | | ERBB2 | | CaMKII | | PIP2 | | ERK | | | |
|---|---|---|---|---|---|---|---|---|---|---|---|---|
| Cluster | Ave | SD | Ave | SD | Ave | SD | Ave | SD | Ave | SD | n | % |
| 1 | -1.02 | 0.32 | -0.70 | 0.22 | -1.10 | 0.41 | -0.81 | 0.33 | -0.88 | 0.37 | 89 | 24.7 |
| 2 | -0.35 | 0.37 | -0.26 | 0.28 | 0.00 | 0.36 | -0.28 | 0.35 | -1.19 | 0.25 | 51 | 14.2 |
| 3 | -0.42 | 0.36 | -0.40 | 0.23 | -0.22 | 0.66 | -0.29 | 0.42 | 0.05 | 0.42 | 96 | 26.7 |
| 4 | 0.15 | 0.35 | 0.03 | 0.34 | 0.34 | 0.47 | 0.12 | 0.47 | 1.67 | 0.56 | 23 | 6.4 |
| 5 | 0.33 | 0.59 | 0.42 | 0.52 | 0.50 | 0.73 | 0.36 | 0.37 | -0.78 | 0.40 | 38 | 10.6 |
| 6 | -0.04 | 0.59 | -0.13 | 0.29 | 0.51 | 0.61 | 1.42 | 0.76 | 0.64 | 0.78 | 34 | 9.4 |
| 7 | 0.98 | 0.36 | 1.99 | 0.65 | 0.92 | 0.57 | 0.83 | 0.65 | -0.46 | 0.63 | 14 | 3.9 |
| 8 | 2.29 | 0.92 | 1.44 | 0.78 | 2.74 | 0.74 | 3.75 | 1.38 | -0.54 | 1.35 | 7 | 1.9 |
| 9 | 2.84 | 0.50 | 4.19 | 0.72 | 2.12 | 0.71 | 1.21 | 0.80 | -0.83 | 0.60 | 8 | 2.2 |

MCF7 + HRG

| | ERBB1 | | ERBB2 | | CaMKII | | PIP2 | | ERK | | | |
|---|---|---|---|---|---|---|---|---|---|---|---|---|
| Cluster | Ave | SD | Ave | SD | Ave | SD | Ave | SD | Ave | SD | n | % |
| 1 | -0.67 | 0.33 | -0.62 | 0.15 | -1.00 | 0.43 | -0.83 | 0.25 | -0.47 | 0.41 | 100 | 26.5 |
| 2 | -0.11 | 0.40 | -0.30 | 0.23 | -0.22 | 0.35 | -0.43 | 0.35 | 0.40 | 0.48 | 88 | 23.3 |
| 3 | -0.37 | 0.44 | -0.36 | 0.27 | -0.19 | 0.62 | 1.20 | 0.69 | 0.17 | 0.72 | 29 | 7.7 |
| 4 | 0.45 | 0.47 | 0.29 | 0.47 | 1.95 | 0.80 | 2.18 | 0.98 | 0.25 | 0.87 | 13 | 3.4 |
| 5 | 0.30 | 0.49 | 0.77 | 0.45 | 0.06 | 0.39 | 0.01 | 0.56 | -0.43 | 0.49 | 22 | 5.8 |
| 6 | 0.78 | 0.73 | 0.05 | 0.38 | 0.72 | 0.44 | 0.05 | 0.52 | 0.75 | 0.48 | 59 | 15.6 |
| 7 | 0.88 | 0.71 | 0.34 | 0.45 | 0.99 | 0.83 | 0.49 | 0.83 | 2.18 | 0.77 | 32 | 8.5 |
| 8 | 1.80 | 0.66 | 2.07 | 0.46 | 1.05 | 0.47 | 0.46 | 0.71 | 0.11 | 0.50 | 22 | 5.8 |
| 9 | 3.64 | 0.84 | 4.38 | 0.93 | 2.07 | 0.80 | 2.25 | 1.57 | 0.33 | 0.84 | 10 | 2.6 |
| 10 | 2.09 | 1.39 | 1.83 | 1.84 | 2.05 | 0.16 | 2.67 | 0.15 | 4.39 | 1.37 | 3 | 0.8 |

### B. ERBB1/B3

HeLa + EGF

| Cluster | ERBB1 Ave | ERBB1 SD | ERBB3 Ave | ERBB3 SD | CaMKII Ave | CaMKII SD | PIP2 Ave | PIP2 SD | ERK Ave | ERK SD | n | % |
|---|---|---|---|---|---|---|---|---|---|---|---|---|
| 1 | -0.40 | 0.31 | -0.63 | 0.34 | -1.09 | 0.42 | -0.62 | 0.52 | 0.15 | 0.40 | 108 | 29.9 |
| 2 | 0.03 | 0.36 | -0.23 | 0.41 | -0.04 | 0.59 | -0.51 | 0.25 | 0.41 | 0.45 | 83 | 23.0 |
| 3 | 0.01 | 0.46 | -0.18 | 0.33 | 0.31 | 0.66 | 1.14 | 0.82 | 0.44 | 0.55 | 23 | 6.4 |
| 4 | 0.76 | 0.58 | 0.64 | 0.42 | -0.06 | 0.42 | -0.55 | 0.31 | 0.85 | 0.52 | 63 | 17.5 |
| 5 | 1.01 | 0.47 | 0.86 | 0.31 | 2.30 | 0.66 | 0.51 | 0.51 | 2.26 | 0.58 | 13 | 3.6 |
| 6 | 1.29 | 0.61 | 1.29 | 0.63 | 0.93 | 0.46 | 0.21 | 0.84 | 0.95 | 0.63 | 54 | 15.0 |
| 7 | 3.49 | 0.81 | 3.38 | 1.01 | 2.10 | 1.07 | 0.29 | 0.69 | 1.54 | 0.93 | 17 | 4.7 |

HeLa + HRG

| Cluster | ERBB1 Ave | ERBB1 SD | ERBB3 Ave | ERBB3 SD | CaMKII Ave | CaMKII SD | PIP2 Ave | PIP2 SD | ERK Ave | ERK SD | n | % |
|---|---|---|---|---|---|---|---|---|---|---|---|---|
| 1 | -0.78 | 0.15 | -0.71 | 0.33 | -0.52 | 0.49 | -0.62 | 0.29 | -1.14 | 0.12 | 65 | 31.9 |
| 2 | -0.76 | 0.15 | -0.59 | 0.34 | -0.13 | 0.54 | 0.47 | 0.52 | -1.10 | 0.13 | 69 | 33.8 |
| 3 | -0.79 | 0.17 | -0.68 | 0.41 | -0.28 | 0.50 | 2.57 | 0.70 | -1.13 | 0.12 | 23 | 11.3 |
| 4 | -0.62 | 0.21 | 0.30 | 0.55 | 0.92 | 0.49 | 0.21 | 0.51 | -0.97 | 0.15 | 33 | 16.2 |
| 5 | -0.58 | 0.27 | 0.40 | 1.03 | 1.52 | 0.71 | 2.60 | 0.85 | -0.85 | 0.30 | 14 | 6.9 |

A431 + EGF

| Cluster | ERBB1 Ave | ERBB1 SD | ERBB3 Ave | ERBB3 SD | CaMKII Ave | CaMKII SD | PIP2 Ave | PIP2 SD | ERK Ave | ERK SD | n | % |
|---|---|---|---|---|---|---|---|---|---|---|---|---|
| 1 | -0.06 | 0.19 | -0.14 | 0.24 | -0.77 | 0.35 | -0.52 | 0.25 | 0.46 | 0.63 | 111 | 29.5 |
| 2 | 0.30 | 0.38 | 0.30 | 0.37 | -0.23 | 0.48 | -0.18 | 0.44 | 0.38 | 0.33 | 106 | 28.2 |
| 3 | 0.23 | 0.34 | 0.22 | 0.42 | 0.26 | 0.28 | 0.07 | 0.50 | 1.68 | 0.50 | 44 | 11.7 |
| 4 | 0.43 | 0.54 | 0.47 | 0.54 | 1.46 | 0.73 | 0.65 | 0.34 | 3.28 | 0.90 | 13 | 3.5 |
| 5 | 0.70 | 0.62 | 0.77 | 0.67 | 1.38 | 0.56 | 0.75 | 0.85 | 1.34 | 0.61 | 28 | 7.4 |
| 6 | 1.29 | 0.41 | 1.16 | 0.62 | -0.28 | 0.76 | -0.16 | 0.62 | -0.49 | 0.47 | 36 | 9.6 |
| 7 | 1.86 | 0.49 | 2.25 | 1.24 | 2.31 | 0.90 | 5.41 | 2.04 | 0.87 | 1.19 | 8 | 2.1 |
| 8 | 2.91 | 0.67 | 2.27 | 0.64 | 0.16 | 0.77 | -0.08 | 0.35 | -0.43 | 0.46 | 24 | 6.4 |
| 9 | 3.47 | 1.66 | 4.76 | 2.40 | 5.02 | 1.07 | 4.19 | 0.08 | 4.35 | 2.21 | 2 | 0.5 |
| 10 | 6.62 | 1.79 | 7.14 | 2.65 | 1.88 | 1.52 | 1.93 | 1.47 | -0.01 | 0.78 | 4 | 1.1 |

A431 + HRG

| Cluster | ERBB1 Ave | ERBB1 SD | ERBB3 Ave | ERBB3 SD | CaMKII Ave | CaMKII SD | PIP2 Ave | PIP2 SD | ERK Ave | ERK SD | n | % |
|---|---|---|---|---|---|---|---|---|---|---|---|---|
| 1 | -0.66 | 0.04 | -0.65 | 0.13 | -1.07 | 0.29 | -0.60 | 0.21 | -0.86 | 0.21 | 68 | 19.0 |
| 2 | -0.63 | 0.05 | -0.57 | 0.19 | -0.14 | 0.38 | -0.23 | 0.31 | -0.63 | 0.34 | 203 | 56.9 |
| 3 | -0.62 | 0.07 | -0.54 | 0.23 | 1.04 | 0.60 | 0.54 | 0.56 | -0.61 | 0.37 | 74 | 20.7 |
| 4 | -0.62 | 0.04 | -0.39 | 0.19 | 3.08 | 1.45 | 3.40 | 1.87 | -0.05 | 0.84 | 12 | 3.4 |

MCF7 + EGF

| Cluster | ERBB1 Ave | ERBB1 SD | ERBB3 Ave | ERBB3 SD | CaMKII Ave | CaMKII SD | PIP2 Ave | PIP2 SD | ERK Ave | ERK SD | n | % |
|---|---|---|---|---|---|---|---|---|---|---|---|---|
| 1 | -1.10 | 0.42 | -1.03 | 0.16 | -1.05 | 0.40 | -0.69 | 0.27 | -0.82 | 0.39 | 81 | 23.3 |
| 2 | -0.59 | 0.53 | -0.81 | 0.25 | -0.16 | 0.57 | 0.29 | 0.75 | 0.25 | 0.57 | 67 | 19.3 |
| 3 | -0.39 | 0.45 | -0.70 | 0.27 | 0.05 | 0.47 | -0.22 | 0.43 | -0.77 | 0.45 | 88 | 25.3 |
| 4 | -0.06 | 0.77 | -0.53 | 0.39 | 0.76 | 0.65 | 1.74 | 0.98 | 1.46 | 1.03 | 30 | 8.6 |
| 5 | -0.03 | 0.72 | -0.65 | 0.28 | 1.77 | 0.66 | 1.47 | 0.76 | -0.91 | 0.49 | 26 | 7.5 |
| 6 | 0.62 | 0.66 | -0.40 | 0.33 | 0.87 | 0.70 | -0.09 | 0.28 | -0.11 | 0.89 | 42 | 12.1 |
| 7 | 1.92 | 0.87 | 0.08 | 0.43 | 2.97 | 1.06 | 4.31 | 0.68 | 2.14 | 0.24 | 4 | 1.1 |
| 8 | 2.39 | 0.74 | -0.22 | 0.38 | 2.14 | 0.77 | 1.25 | 0.68 | 0.28 | 0.92 | 10 | 2.9 |

MCF7 + HRG

| Cluster | ERBB1 Ave | ERBB1 SD | ERBB3 Ave | ERBB3 SD | CaMKII Ave | CaMKII SD | PIP2 Ave | PIP2 SD | ERK Ave | ERK SD | n | % |
|---|---|---|---|---|---|---|---|---|---|---|---|---|
| 1 | -0.53 | 0.45 | 0.06 | 0.43 | -1.12 | 0.41 | -0.81 | 0.24 | -0.51 | 0.43 | 98 | 26.3 |
| 2 | -0.35 | 0.48 | 0.02 | 0.39 | -0.18 | 0.46 | 0.09 | 0.78 | 0.67 | 0.58 | 54 | 14.5 |
| 3 | 0.42 | 0.80 | -0.44 | 0.34 | 1.11 | 0.68 | 1.73 | 1.60 | -1.28 | 0.11 | 9 | 2.4 |
| 4 | 0.67 | 0.60 | 0.37 | 0.45 | 1.17 | 0.58 | 2.21 | 0.85 | 1.03 | 0.66 | 18 | 4.8 |
| 5 | 0.73 | 0.49 | 0.55 | 0.52 | 0.76 | 0.46 | 0.00 | 0.33 | 0.35 | 0.39 | 27 | 7.3 |
| 6 | 0.54 | 0.68 | 0.78 | 0.55 | 0.77 | 0.68 | 0.30 | 0.77 | 2.36 | 0.93 | 29 | 7.8 |
| 7 | 0.53 | 0.84 | 1.11 | 0.62 | -0.27 | 0.45 | -0.46 | 0.49 | -0.09 | 0.39 | 98 | 26.3 |
| 8 | 1.95 | 0.64 | 2.41 | 0.89 | 0.64 | 0.69 | -0.15 | 0.36 | 0.72 | 0.67 | 39 | 10.5 |

### C. ERBB1/B4

HeLa + EGF

| Cluster | ERBB1 Ave | ERBB1 SD | ERBB4 Ave | ERBB4 SD | CaMKII Ave | CaMKII SD | PIP2 Ave | PIP2 SD | ERK Ave | ERK SD | n | % |
|---|---|---|---|---|---|---|---|---|---|---|---|---|
| 1 | -0.53 | 0.20 | -0.42 | 0.26 | -1.35 | 0.35 | -0.79 | 0.30 | 0.37 | 0.47 | 46 | 12.2 |
| 2 | 0.31 | 0.34 | 0.29 | 0.31 | -0.66 | 0.44 | -0.62 | 0.28 | 0.43 | 0.29 | 71 | 18.8 |
| 3 | -0.35 | 0.25 | -0.42 | 0.22 | -0.28 | 0.41 | -0.44 | 0.42 | 0.21 | 0.36 | 73 | 19.4 |
| 4 | 0.38 | 0.55 | 0.28 | 0.48 | 0.49 | 0.56 | -0.38 | 0.25 | 1.21 | 0.50 | 82 | 21.8 |
| 5 | 0.09 | 0.73 | 0.14 | 0.71 | 0.35 | 0.92 | 1.49 | 0.55 | 0.37 | 0.57 | 10 | 2.7 |
| 6 | 1.53 | 0.52 | 1.64 | 0.61 | 0.12 | 0.60 | -0.36 | 0.47 | 0.72 | 0.59 | 68 | 18.0 |
| 7 | 1.76 | 0.78 | 1.71 | 0.63 | 2.39 | 0.86 | 0.13 | 0.43 | 1.88 | 0.53 | 12 | 3.2 |
| 8 | 3.95 | 0.97 | 3.83 | 0.59 | 1.12 | 0.82 | 0.05 | 0.30 | 1.71 | 1.03 | 8 | 2.1 |
| 9 | 3.54 | 1.06 | 3.59 | 0.92 | 3.13 | 0.71 | 2.79 | 1.30 | 3.01 | 0.66 | 7 | 1.9 |

HeLa + HRG

| Cluster | ERBB1 Ave | ERBB1 SD | ERBB4 Ave | ERBB4 SD | CaMKII Ave | CaMKII SD | PIP2 Ave | PIP2 SD | ERK Ave | ERK SD | n | % |
|---|---|---|---|---|---|---|---|---|---|---|---|---|
| 1 | -0.70 | 0.18 | -0.72 | 0.20 | -0.48 | 0.53 | -0.27 | 0.46 | -1.00 | 0.22 | 153 | 52.8 |
| 2 | -0.72 | 0.13 | -0.71 | 0.15 | -0.32 | 0.35 | 1.32 | 0.60 | -1.02 | 0.17 | 45 | 15.5 |
| 3 | -0.56 | 0.24 | -0.43 | 0.25 | 1.12 | 0.85 | 1.80 | 0.52 | -0.81 | 0.21 | 40 | 13.8 |
| 4 | -0.61 | 0.19 | -0.45 | 0.30 | 1.19 | 1.57 | 4.01 | 0.72 | -0.79 | 0.38 | 9 | 3.1 |
| 5 | -0.42 | 0.21 | -0.45 | 0.19 | 1.18 | 0.54 | 0.12 | 0.49 | -0.78 | 0.24 | 43 | 14.8 |

A431 + EGF

| Cluster | ERBB1 Ave | ERBB1 SD | ERBB4 Ave | ERBB4 SD | CaMKII Ave | CaMKII SD | PIP2 Ave | PIP2 SD | ERK Ave | ERK SD | n | % |
|---|---|---|---|---|---|---|---|---|---|---|---|---|
| 1 | -0.10 | 0.17 | -0.07 | 0.23 | -0.69 | 0.30 | -0.48 | 0.21 | 0.36 | 0.43 | 87 | 24.9 |
| 2 | -0.09 | 0.23 | -0.03 | 0.34 | -0.35 | 0.42 | -0.33 | 0.21 | 1.62 | 0.57 | 33 | 9.5 |
| 3 | 0.47 | 0.54 | 0.64 | 0.54 | -0.34 | 0.57 | -0.05 | 1.34 | 0.20 | 0.41 | 81 | 23.2 |
| 4 | 0.67 | 0.39 | 0.84 | 0.32 | 0.42 | 0.54 | 0.30 | 0.65 | 0.95 | 0.57 | 60 | 17.2 |
| 5 | 0.67 | 0.70 | 0.88 | 0.63 | 0.90 | 0.47 | 0.29 | 0.50 | 2.57 | 1.09 | 24 | 6.9 |
| 6 | 1.48 | 0.50 | 1.60 | 0.43 | -0.40 | 0.46 | -0.14 | 0.39 | -0.35 | 0.49 | 38 | 10.9 |
| 7 | 3.95 | 1.28 | 3.25 | 0.83 | 0.23 | 0.70 | 0.18 | 0.53 | -0.57 | 0.30 | 12 | 3.4 |
| 8 | 2.45 | 0.63 | 2.34 | 0.41 | 2.07 | 1.00 | 1.88 | 0.80 | 0.77 | 1.22 | 9 | 2.6 |
| 9 | 3.61 | 2.05 | 3.44 | 0.95 | 4.65 | 0.85 | 4.39 | 1.52 | 3.49 | 1.35 | 5 | 1.4 |

A431 + HRG

| Cluster | ERBB1 Ave | ERBB1 SD | ERBB4 Ave | ERBB4 SD | CaMKII Ave | CaMKII SD | PIP2 Ave | PIP2 SD | ERK Ave | ERK SD | n | % |
|---|---|---|---|---|---|---|---|---|---|---|---|---|
| 1 | -0.67 | 0.07 | -0.76 | 0.15 | -0.71 | 0.26 | -0.49 | 0.23 | -0.76 | 0.19 | 144 | 41.6 |
| 2 | -0.65 | 0.08 | -0.73 | 0.19 | 0.09 | 0.39 | -0.08 | 0.33 | -0.60 | 0.39 | 144 | 41.6 |
| 3 | -0.65 | 0.08 | -0.71 | 0.18 | 1.63 | 0.61 | 0.48 | 0.64 | -0.45 | 0.52 | 42 | 12.1 |
| 4 | -0.64 | 0.08 | -0.64 | 0.25 | 2.70 | 1.18 | 3.67 | 1.36 | -0.48 | 0.44 | 16 | 4.6 |

MCF7 + EGF

| Cluster | ERBB1 Ave | ERBB1 SD | ERBB4 Ave | ERBB4 SD | CaMKII Ave | CaMKII SD | PIP2 Ave | PIP2 SD | ERK Ave | ERK SD | n | % |
|---|---|---|---|---|---|---|---|---|---|---|---|---|
| 1 | -0.93 | 0.50 | -0.95 | 0.29 | -1.04 | 0.35 | -0.72 | 0.31 | -0.73 | 0.37 | 109 | 30.6 |
| 2 | -0.59 | 0.38 | -0.71 | 0.37 | -0.28 | 0.37 | -0.25 | 0.40 | -1.04 | 0.40 | 57 | 16.0 |
| 3 | -0.44 | 0.45 | -0.54 | 0.30 | -0.49 | 0.41 | -0.22 | 0.71 | 0.42 | 0.65 | 66 | 18.5 |
| 4 | 0.13 | 0.45 | -0.14 | 0.39 | 0.59 | 0.45 | 0.07 | 0.44 | -0.61 | 0.60 | 53 | 14.9 |
| 5 | -0.43 | 0.57 | 0.01 | 0.40 | 1.22 | 0.43 | 1.79 | 0.52 | -0.62 | 0.65 | 12 | 3.4 |
| 6 | 0.53 | 0.97 | 0.15 | 0.61 | 1.65 | 0.76 | 4.06 | 1.19 | 0.41 | 1.09 | 9 | 2.5 |
| 7 | 0.83 | 0.53 | 0.25 | 0.53 | 0.50 | 0.41 | 0.11 | 0.38 | 1.17 | 0.84 | 23 | 6.5 |
| 8 | 0.99 | 0.61 | 0.86 | 0.72 | 1.97 | 0.71 | 0.80 | 0.51 | 0.44 | 0.94 | 21 | 5.9 |
| 9 | 1.10 | 1.14 | 0.47 | 0.77 | 1.99 | 0.23 | 1.31 | 0.98 | 3.15 | 0.92 | 6 | 1.7 |

MCF7 + HRG

| Cluster | ERBB1 Ave | ERBB1 SD | ERBB4 Ave | ERBB4 SD | CaMKII Ave | CaMKII SD | PIP2 Ave | PIP2 SD | ERK Ave | ERK SD | n | % |
|---|---|---|---|---|---|---|---|---|---|---|---|---|
| 1 | -0.52 | 0.49 | -0.41 | 0.37 | -1.00 | 0.31 | -0.67 | 0.21 | -0.54 | 0.37 | 88 | 23.7 |
| 2 | -0.44 | 0.41 | -0.30 | 0.36 | -0.30 | 0.39 | -0.06 | 0.61 | 0.24 | 0.41 | 54 | 14.5 |
| 3 | 0.33 | 0.31 | 0.33 | 0.41 | -0.03 | 0.46 | -0.28 | 0.26 | 0.06 | 0.58 | 86 | 23.1 |
| 4 | -0.11 | 0.67 | 0.35 | 0.68 | 0.60 | 0.45 | 0.42 | 0.64 | 1.77 | 0.78 | 27 | 7.3 |
| 5 | 0.33 | 0.60 | 0.33 | 0.73 | 1.18 | 0.59 | 1.77 | 0.84 | 0.07 | 0.85 | 17 | 4.6 |
| 6 | -0.06 | 0.36 | -0.31 | 0.26 | 1.57 | 0.64 | 4.77 | 0.93 | -1.03 | 0.35 | 4 | 1.1 |
| 7 | 1.18 | 0.62 | 1.35 | 0.62 | 0.89 | 0.75 | 0.11 | 0.43 | 0.53 | 0.76 | 70 | 18.8 |
| 8 | 1.26 | 0.70 | 1.81 | 1.33 | 2.16 | 1.03 | 2.76 | 0.94 | 2.36 | 1.04 | 9 | 2.4 |
| 9 | 3.05 | 0.91 | 3.48 | 1.14 | 1.16 | 0.50 | 0.45 | 0.45 | 1.12 | 1.05 | 16 | 4.3 |
| 10 | 7.45 | NA | 2.42 | NA | -0.15 | NA | 3.95 | NA | 0.64 | NA | 1 | 0.3 |
